# SNPC-1.3 is a sex-specific transcription factor that drives male piRNA expression in *C. elegans*

**DOI:** 10.1101/2020.08.06.240200

**Authors:** Charlotte P. Choi, Rebecca J. Tay, Margaret R. Starostik, Suhua Feng, James J. Moresco, Brooke E. Montgomery, Emily Xu, Maya A. Hammonds, Michael C. Schatz, Taiowa A. Montgomery, John R. Yates, Steven E. Jacobsen, John K. Kim

## Abstract

Piwi-interacting RNAs (piRNAs) play essential roles in silencing repetitive elements to promote fertility in metazoans. Studies in worms, flies, and mammals reveal that piRNAs are expressed in a sex-specific manner. However, the mechanisms underlying this sex-specific regulation are unknown. Here we identify SNPC-1.3, a variant of a conserved subunit of the snRNA activating protein complex, as a male-specific piRNA transcription factor in *C. elegans*. Binding of SNPC-1.3 at male piRNA loci drives spermatogenic piRNA transcription and requires the core piRNA transcription factor SNPC-4. Loss of *snpc-1.3* leads to depletion of male piRNAs and defects in male-dependent fertility. Furthermore, TRA-1, a master regulator of sex determination, binds to the *snpc-1.3* promoter and represses its expression during oogenesis. Loss of TRA-1 targeting causes ectopic expression of *snpc-1.3* and male piRNAs during oogenesis. Thus, sexual dimorphic regulation of *snpc-1.3* coordinates male and female piRNA expression during germline development.

## INTRODUCTION

Piwi-interacting RNAs (piRNAs), a distinct class of small noncoding RNAs, function to preserve germline integrity (Batista et al., 2008; Carmell et al., 2007; Cox et al., 1998; Deng and Lin, 2002; Kuramochi-Miyagawa et al., 2008; Lin and Spradling, 1997; Wang and Reinke, 2008). In *Drosophila*, mutation of any of the three Piwi genes (*piwi*, *aub*, *ago3*) results in rampant activation of transposons in the germline and severe defects in fertility (Brennecke et al., 2007; Harris and Macdonald, 2001; Lin and Spradling, 1997; Vagin et al., 2006). In *M. musculus*, mutation of the Piwi protein *Miwi* leads to the misregulation of genes involved in germ cell development, defective gametogenesis, and sterility (Deng and Lin, 2002; Zhang et al., 2015b). *C. elegans* piRNAs can be inherited across multiple generations and trigger the transgenerational silencing of coding genes. Disruption of this inheritance results in eventual germline collapse and sterility, known as the germline mortal phenotype (Ashe et al., 2012; Buckley et al., 2012; Shirayama et al., 2012). Taken together, piRNAs are essential to preserve germline integrity and protect the reproductive capacity in metazoans.

Loss of the piRNA pathway can have distinct consequences between sexes and across developmental stages. Many species show sex-specific expression of piRNAs (Armisen et al., 2009; Billi et al., 2013; Williams et al., 2015; Yang et al., 2013; Zhou et al., 2010). Demonstrated by hybrid dysgenesis, the identity of female, but not male, piRNAs in flies is important for fertility (Brennecke et al., 2008). In contrast, the piRNA pathway in mammals appears to be dispensable for female fertility (Carmell et al., 2007; Murchison et al., 2007), but distinct subsets of piRNAs are required for specific stages of spermatogenesis (Aravin et al., 2003; Aravin et al., 2006; Carmell et al., 2007; Di Giacomo et al., 2013; Gainetdinov et al., 2018; Girard et al., 2006; Grivna et al., 2006; Kuramochi-Miyagawa et al., 2008; Li et al., 2013). In worms, most piRNAs are uniquely enriched in either the male or female germline (Billi et al., 2013; Kato et al., 2009). Nevertheless, in all of these contexts, how the specific expression of different piRNA subclasses is achieved is poorly understood.

piRNA biogenesis is strikingly diverse across organisms and tissue types. In the *Drosophila* germline, piRNA clusters are found within pericentromeric or telomeric heterochromatin enriched for H3K9me3 histone modifications. The HP1 homolog Rhino binds to H3K9me3 within most of these piRNA clusters and recruits Moonshiner, a paralog of the basal transcription factor TFIIA, which, in turn, recruits RNA polymerase II (Pol II) to enable transcription within heterochromatin (Andersen et al., 2017; Chen et al., 2016; Klattenhoff et al., 2009; Mohn 2014; et al., Pane et al., 2011). Two waves of piRNA expression occur in mouse testes: pre-pachytene piRNAs are expressed in early spermatogenesis and silence transposons, whereas pachytene piRNAs are expressed in the later stages of meiosis and have unknown functions. While the mechanisms of pre-pachytene piRNA transcription remain elusive, pachytene piRNAs require the transcription factor A-MYB, along with RNA Pol II (Li et al., 2013).

In *C. elegans*, SNPC-4 is essential for the expression of piRNAs in the germline (Kasper et al., 2014). SNPC-4 is the single *C. elegans* ortholog of mammalian SNAPC4, the largest DNA binding subunit of the small nuclear RNA (snRNA) activating protein complex (SNAPc). A complex of SNAPC4, SNAPC1, and SNAPC3 binds to the proximal sequence element (PSE) of snRNAs to promote their transcription (Henry et al., 1995; Jawdekar and Henry, 2008; Ma and Hernandez, 2002; Su et al., 1997; Wong et al., 1998; Yoon et al., 1995). SNPC-4 occupies transcription start sites of other classes of noncoding RNAs across various *C. elegans* tissue types and developmental stages (Kasper et al., 2014; Weng et al., 2018). Furthermore, piRNA biogenesis factors PRDE-1, TOFU-4, and TOFU-5 are expressed in germ cell nuclei and interact with SNPC-4 at clusters of piRNA loci (Goh et al., 2014; Kasper et al., 2014; Weick et al., 2014; Weng et al., 2018). These data suggest that SNPC-4 has been co-opted by germline-specific factors to transcribe piRNAs.

The vast majority of the ~15,000 piRNAs in *C. elegans* are encoded within two large megabase genomic clusters on chromosome IV (Das et al., 2008; Ruby et al., 2006). Each piRNA locus encodes a discrete transcriptional unit that is individually transcribed as a short precursor by Pol II (Gu et al., 2012; Cecere et al., 2012; Billi et al., 2013). Processing of precursors yields mature piRNAs that are typically 21 nucleotides (nt) in length and strongly enriched for a 5’ uracil (referred to as 21U-RNAs). Transcription of these piRNAs requires a conserved 8 nt core motif (NNGTTTCA) within their promoters (Billi et al., 2013; Cecere et al., 2012; Ruby et al., 2006). piRNAs enriched during spermatogenesis are associated with a cytosine at the 5’ most position of the core motif (<u>C</u>NGTTTCA); mutation of cytosine to adenine at this position results in ectopic expression of normally male-enriched piRNAs during oogenesis. In contrast, genomic loci expressing piRNAs enriched in the female germline show no discernable nucleotide bias at the 5’ position. While differences in *cis*-regulatory sequences contribute to the sexually dimorphic nature of piRNA expression, sex-specific piRNA transcription factors that drive distinct subsets of piRNAs in the male and female germlines remain to be identified.

Here, we discovered that SNPC-1.3, an ortholog of human SNAPC1, is required specifically for male piRNA expression. Furthermore, TRA-1, a master regulator of sex-determination, transcriptionally represses *snpc-1.3* during oogenesis to restrict its expression to the male germline. Taken together, our study reports the first example of a sex-specific piRNA transcription factor that drives the expression of male-specific piRNAs.

## RESULTS

### SNPC-4 is a component of the core piRNA transcription complex that drives all piRNA expression

SNPC-4-specific foci are present in both male and female germ cell nuclei (Kasper et al., 2014), but the role of SNPC-4 in the male germline is not well understood. We hypothesized that SNPC-4 is required for piRNA biogenesis in both the male and female germlines. To test this, we conditionally depleted the SNPC-4 protein using the auxin-inducible degradation system (Zhang et al., 2015a) (Figure S1A). We added an auxin-inducible degron (AID) to the C-terminus of SNPC-4 using CRISPR/Cas9 genome engineering, and crossed this strain into worms expressing TIR1 under the germline promoter, *sun-1*. TIR1 is a plant-specific F-box protein that mediates the rapid degradation of *C. elegans* proteins tagged with an AID in the presence of the phytohormone auxin. Thus, addition of auxin to the *snpc-4::aid; P_sun-1_::TIR1* strain is expected to degrade SNPC-4::AID, whereas strains with *snpc-4::aid* alone serve as a negative control; under these conditions, we examined a panel of spermatogenesis- and oogenesis-enriched piRNAs (Billi et al., 2013). We found that worms depleted of SNPC-4 showed decreased expression of both spermatogenesis- and oogenesis-enriched piRNAs during spermatogenesis and oogenesis, respectively (Figure 1A), indicating that SNPC-4 is a core piRNA transcription factor required for all piRNA expression.

**Figure 1.**
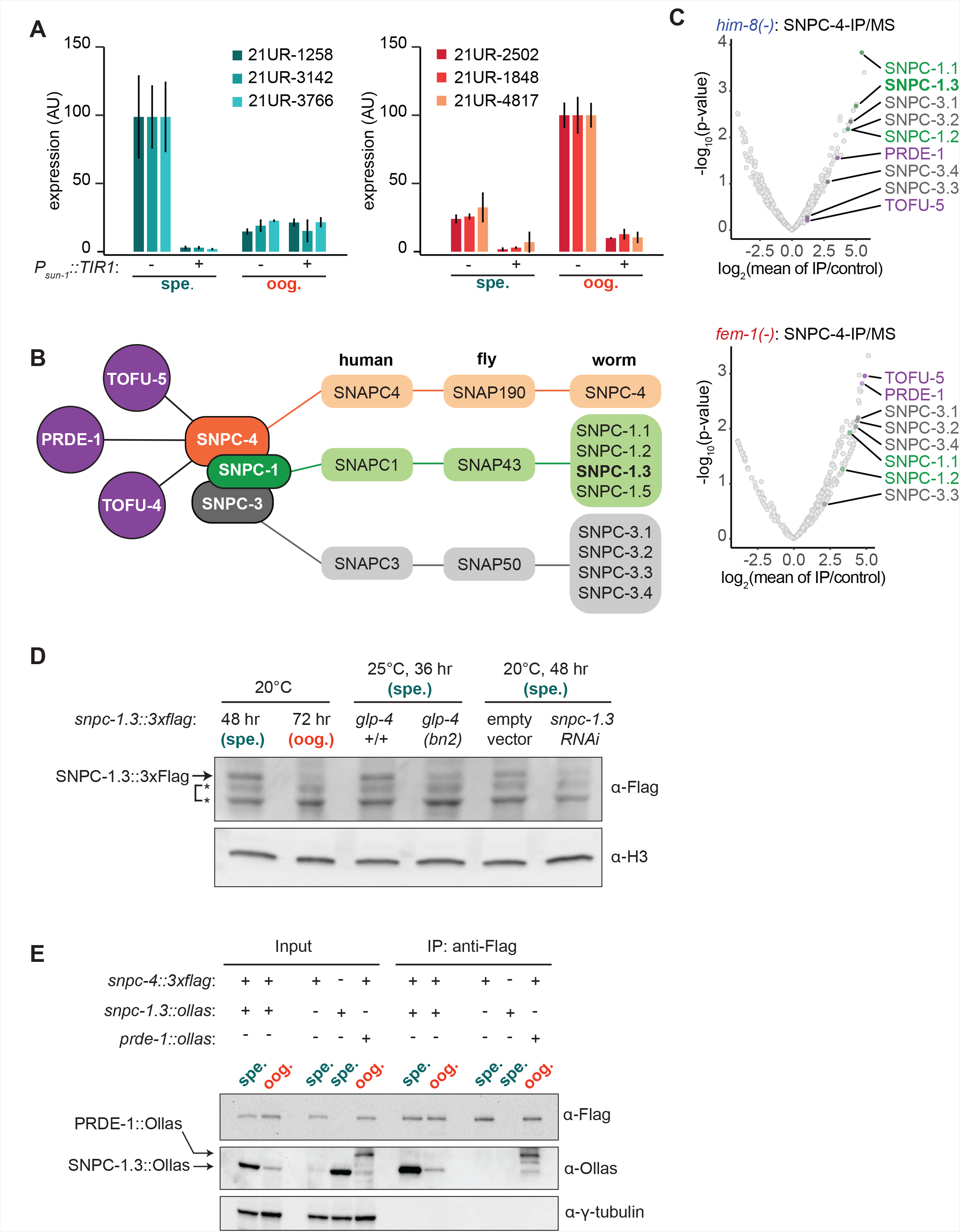
SNPC-4 and SNPC-1.3 are part of the male piRNA transcription complex. (A) SNPC-4 is required for both male and female piRNA expression. Taqman qPCR of male (left) and female (right) piRNAs during spermatogenesis and oogenesis in *snpc-4::aid* (denoted as -) and *snpc-4::aid; P_sun-1_::TIR1* (denoted as +) worms. Error bars indicate ± SD from two technical replicates. (B) Schematic highlights the conservation of SNAPc homologs from *C. elegans, D. melanogaster, and H. sapiens* and catalogs all SNPC-4 (orange) interacting partners from previous work (Weick et al., 2014; Weng et al., 2018) or from our own analysis. Known piRNA biogenesis factors (purple), SNPC-1 paralogs (green), and SNPC-3 paralogs (grey) are indicated. (C) SNPC-1.3 interacts with SNPC-4 in only *him-8(-)* mutants. Volcano plots showing enrichment values and analogous significance values for proteins that co-purified with SNPC-4::3xFlag from (top) *him-8(-)* mutants (n=2 biological replicates) or (bottom) *fem-1(-)* mutants (n=2 biological replicates). piRNA biogenesis factors (purple), SNPC-1 paralogs (green), and SNPC-3 paralogs (dark grey) are labeled in Figure 1B. (D) SNPC-1.3 is predominantly expressed in the male germline. Western blot of SNPC-1.3::3xFlag expression during spermatogenesis (spe.) and oogenesis (oog.) at 20°C and expression in germline-less *glp-4(bn2)* worms at 25°C during spermatogenesis (spe.). RNAi of *snpc-1.3* served to identify the SNPC-1.3::3xFlag band; * denotes non-specific bands. (E) SNPC-4 interacts with SNPC-1.3. Anti-Flag immunoprecipitation of SNPC-4::3xFlag and western blot for SNPC-1.3::Ollas during spermatogenesis (spe.) and oogenesis (oog.). PRDE::Ollas was used as a positive control for interaction with SNPC-4::3xFlag (Kasper et al., 2014). γ-tubulin was used as the loading control.

Given that SNPC-4 activates transcription of piRNAs in both sexes, we hypothesized that sex-specific cofactors might associate with SNPC-4 to regulate sexually dimorphic piRNA expression. To test this hypothesis, we leveraged genetic backgrounds that masculinize or feminize the germline. Specifically, we used *him-8(-)* mutants, which have a higher incidence of males (∼30% males compared to <0.5% spontaneous males in the wild-type hermaphrodite population) (Hodgkin et al., 1979), and *fem-1(-)* mutants, which are completely feminized when grown at 25°C (Doniach and Hodgkin, 1984). We introduced a C-terminal 3xFlag tag sequence at the endogenous *snpc-4* locus using CRISPR/Cas9 genome editing (Paix et al., 2015) and performed immunoprecipitation of SNPC-4::3xFlag followed by mass spectrometry. We identified PRDE-1 and TOFU-5 as co-purifying with SNPC-4::3xFlag in both *him-8(-)* and *fem-1(-)* mutants, suggesting that these known piRNA biogenesis factors exist as a complex in both male and female germlines (Figures 1B–C, and S1B). While a single worm ortholog, SNPC-4, exists for human SNAPC4, the *C. elegans* genome encodes 4 homologs of human SNAPC1 (worm SNPC-1.1, −1.2, - 1.3, and −1.5) and 4 homologs of human SNAPC3 (worm SNPC-3.1, −3.2, −3.3, and −3.4) (Figure 1B) (Li et al., 2004). From our mass spectrometry analysis, 6 out of the 8 *C. elegans* homologs of SNAPC1 and SNAPC3 co-purified with SNPC-4::3xFlag from both *him-8(-)* and *fem-1(-)* genetic backgrounds (Figures 1B–C). These results revealed that SNPC-4 interacts with both snRNA and piRNA transcription machinery.

### SNPC-1.3 interacts with the core piRNA biogenesis factor SNPC-4 during spermatogenesis

We also identified proteins that co-purified with SNPC-4::3xFlag from *him-8(-),* but not *fem-1(-)* mutants. We were particularly interested in SNPC-1.3 because of its homology to the mammalian SNAPC1 subunit of the snRNA transcription complex. To characterize SNPC-1.3, we used CRISPR/Cas9 genome editing to insert a 3xFlag tag sequence at the C-terminus of the endogenous *snpc-1.3* locus. Addition of 3xFlag at the C-terminus of either the a or b isoform had no effect on fertility or levels of spermatogenesis- and oogenesis-enriched piRNAs (Figures S1C–G). Henceforth, SNPC-1.3 refers specifically to the SNPC-1.3a isoform. SNPC-1.3::3xFlag was highly expressed during spermatogenesis and showed markedly reduced expression during oogenesis (Figure 1D). To determine whether SNPC-1.3 expression is restricted to the germline, we examined SNPC-1.3::3xFlag expression in the *glp-4(bn2)* mutant, which fails to develop fully-expanded germlines at 25°C (Beanan and Strome, 1992). SNPC-1.3::3xFlag expression during early spermatogenesis was greatly reduced in *glp-4(bn2)* compared to wildtype, suggesting that SNPC-1.3 is primarily expressed in the germline during spermatogenesis (Figure 1D).

To confirm that SNPC-1.3 interacts with SNPC-4, we used CRISPR/Cas9 genome editing to generate an endogenously tagged *snpc-1.3::ollas* strain. We then crossed *snpc-1.3::ollas* into the *snpc-4::3xflag* strain and performed co-immunoprecipitation experiments with anti-Flag antibodies. In agreement with the mass spectrometry data, SNPC-4::3xFlag and SNPC-1.3:Ollas interacted robustly during spermatogenesis. The interaction was detectable at a much lower level during oogenesis (Figure 1E), likely due to the significant decrease in SNPC-1.3 expression during this time (Figure 1D). The reciprocal co-immunoprecipitation of SNPC-1.3::3xFlag followed by western blotting for SNPC-4::Ollas confirmed this biochemical interaction (Figure S1H). Taken together, our data indicate that SNPC-1.3 forms a complex with the previously characterized piRNA biogenesis factor SNPC-4 in the male germline.

### SNPC-1.3 is required for transcription of male piRNAs

Given the prominent interaction between SNPC-1.3 and SNPC-4 in the male germline (Figure 1E), we hypothesized that SNPC-1.3 might be required for piRNA expression during spermatogenesis. To test this hypothesis, we generated a *snpc-1.3* null allele by introducing mutations that result in a premature stop codon (Paix et al., 2015). We examined spermatogenesis in hermaphrodites and *him-8(-)* males, and examined oogenesis in adult hermaphrodites and *fem-1(-)* females. As a control, we analyzed the loss-of-function mutant of the *C. elegans* Piwi protein, *prg-1(-)*, which almost completely lacked spermatogenesis- and oogenesis-enriched piRNAs (Figure 2A), as expected. We found that the levels of spermatogenesis-enriched piRNAs were dramatically reduced in *snpc-1.3(-)* hermaphrodites during spermatogenesis and in *him-8(-); snpc-1.3(-)* males, whereas oogenesis-enriched piRNAs were largely unaltered in *snpc-1.3(-)* adult hermaphrodites and in *fem-1(-); snpc-1.3(-)* females (Figures 2A–B). Moreover, oogenesis-enriched piRNAs were upregulated at least 2-fold in *snpc-1.3(-)* mutants undergoing spermatogenesis and in *him-8(-); snpc-1.3(-)* males. These findings suggest that, in addition to activating male piRNAs, SNPC-1.3 suppresses the expression of female piRNAs in the male germline, possibly by preferentially recruiting core factors such as SNPC-4 to male piRNA loci.

**Figure 2.**
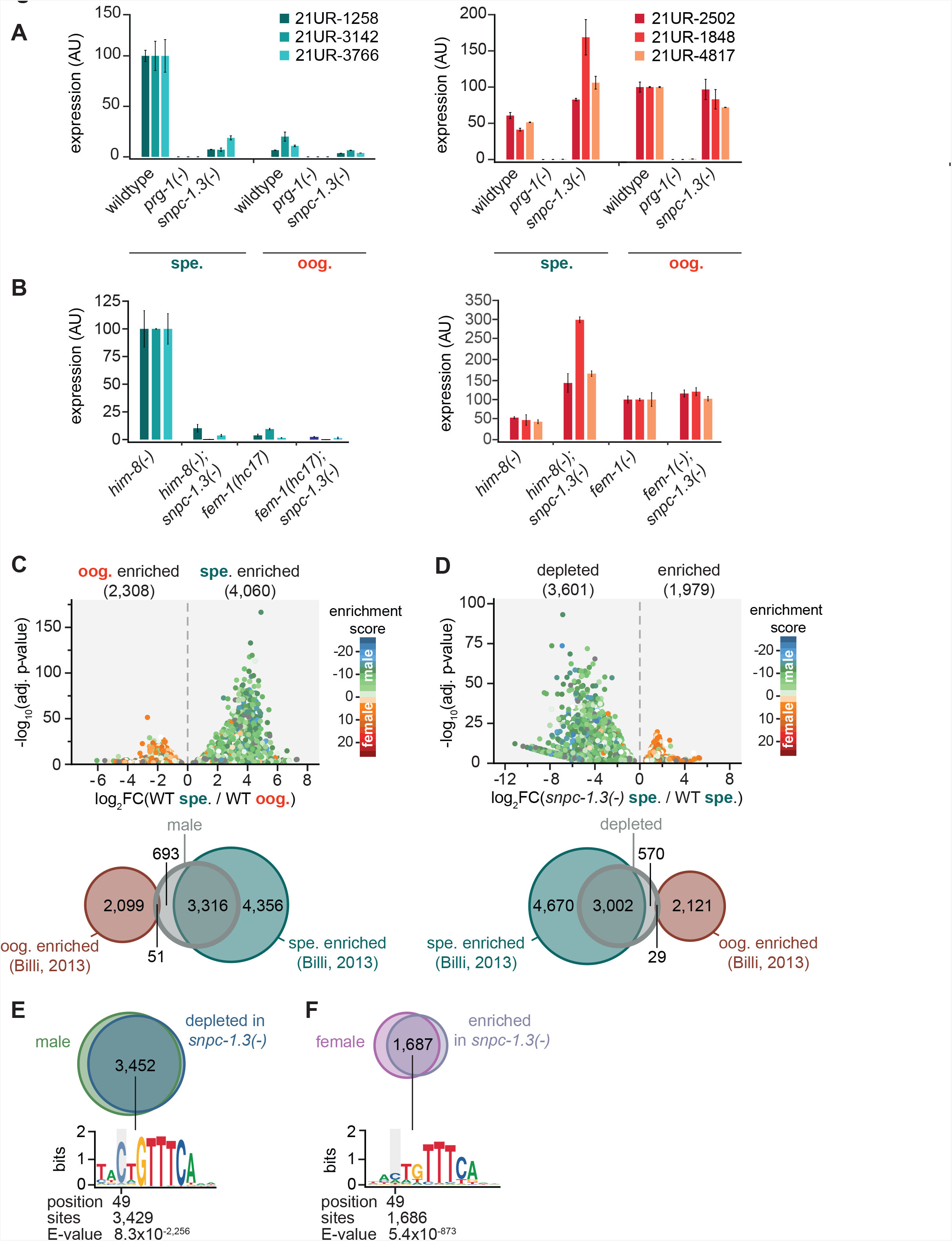
SNPC-1.3 is required for transcription of male piRNAs. (A) *snpc-1.3* is required for male piRNA expression during spermatogenesis (spe.) but is dispensable for female piRNA expression during oogenesis (oog.). Taqman qPCR and quantification of representative male (left) and female (right) piRNAs at spermatogenic and oogenic time points. Error bars indicate ± SD from two technical replicates. (B) *him-8(-)*; *snpc-1.3(-)* mutant males exhibit severely impaired male piRNA expression and enhanced female piRNA expression. *snpc-1.3* is not required for male or female piRNA expression in *fem-1(hc17)* females. Error bars indicate ± SD from two technical replicates. (C) piRNAs are differentially expressed during spermatogenesis (spe.) and oogenesis (oog.) in wild-type worms. (Top) Volcano plot showing piRNAs with ≥1.2-fold change and FDR of ≤ 0.05 in 48 h (spe.) versus 72 h (oog.). piRNAs are colored according to male and female enrichment scores from Billi et al., 2013. (Bottom) Overlap of male piRNAs in wildtype at 48 h (spe.) with oogenesis- and spermatogenesis-enriched piRNAs defined in from Billi et al., 2013. (D) piRNAs depleted in *snpc-1.3(-)* comprise mostly of spermatogenesis-enriched piRNAs. (Top) Volcano plot shows piRNAs with ≥1.2-fold change and FDR < 0.05 in *snpc-1.3(-)* mutant versus wildtype during spermatogenesis (spe.). piRNAs are colored according to male and female enrichment scores from Billi et al., 2013. (Bottom) Overlap of *snpc-1.3-*dependent piRNAs with oogenesis (oog.)- and spermatogenesis (spe.)-enriched piRNAs defined in Billi et al., 2013. (E) Male piRNAs that are depleted *snpc-1.3(-)* have a conserved upstream motif with a strong 5’ C bias. (Top) Overlap of *snpc-1.3-*dependent piRNAs with male piRNAs shown in Figure 2C. (Bottom) Logo plot displays conserved motif upstream of each piRNA. Median position of the C-nucleotide of the identified motif, number of piRNAs sharing the motif, and associated E-value are listed. (F) Female piRNAs are upregulated in *snpc-1.3(-)* mutants during spermatogenesis. (Top) Overlap of piRNAs upregulated at 72 h (oog.) with piRNAs enriched in *snpc-1.3(-)* at 48 h (spe.). (Bottom) Logo plot displays conserved motif upstream of each piRNA. Median position of the C-nucleotide of the identified motif, number of piRNAs sharing the motif, and associated E-value are listed.

To extend these findings, we identified piRNAs enriched during spermatogenesis and oogenesis by small RNA-seq in wild-type worms (Figures S2A and S3A). Using a 1.2-fold threshold and false discovery rate (FDR) of ≤ 0.05, a total of 6,368 out of 14,714 piRNAs on chromosome IV were differentially expressed (Figures 2C, S2, S3B, and Table S1). Among these, 4,060 piRNAs were upregulated during spermatogenesis (hereafter referred to as male piRNAs) and 2,308 piRNAs were upregulated during oogenesis, which we define as female piRNAs. We compared this dataset with our previous study that identified spermatogenesis- and oogenesis-enriched piRNAs (Billi et al., 2013). Most male piRNAs identified here were also found in our previous study (82%; 3,316/4,060) (Figure 2C). Next, we investigated how loss of *snpc-1.3* affects global piRNA expression by performing small RNA-seq in wildtype versus *snpc-1.3(-)* mutants during spermatogenesis. We identified 3,601 piRNAs that were downregulated in a *snpc-1.3(-)* mutant compared to wildtype (Figures 2D, S3C, and Table S2). Of these, 3,002 overlapped with spermatogenesis-enriched piRNAs identified in our previous study (Billi et al., 2013) (Figure 2D). 85% (3,452/4,060) of male piRNAs were also depleted in *snpc-1.3(-)* mutants during spermatogenesis, suggesting that male piRNAs are regulated by SNPC-1.3 (Figures 2E and S3D). In addition, 73% (1,687/2,308) of female piRNAs were ectopically upregulated in *snpc-1.3(-)* mutants during spermatogenesis (Figure 2F).

We next analyzed the genomic loci of male piRNAs and *snpc-1.3*-dependent piRNAs. As expected, the intersection of these two piRNA subsets displayed strong enrichment for the 8 nt core motif and the 5’-most position of this core motif was enriched for cytosine (<u>C</u>NGTTTCA) (Figures 2E and S3D). In contrast, the core motif found upstream of oogenesis-enriched piRNAs upregulated upon loss of *snpc-1.3* displayed a much weaker bias for the 5’ cytosine (Figure 2F). These observations validate our previous findings that male and female core motifs are distinct (Billi et al., 2013). Taken together, these data indicate that SNPC-1.3 is required for male piRNA expression.

### SNPC-1.3 binds male piRNA loci in a SNPC-4-dependent manner

Given that SNPC-1.3 interacts with SNPC-4 and is required for expression of male piRNAs, we hypothesized that SNPC-1.3 might bind male piRNA loci in association with SNPC-4. To test this, we performed ChIP-qPCR to investigate SNPC-1.3 occupancy at regions of high piRNA density within the two large piRNA clusters on chromosome IV; an intergenic region lacking piRNAs served as a control. To determine whether SNPC-1.3 binding was dependent on SNPC-4, we again used the auxin-inducible degradation system to deplete SNPC-4 in the *snpc-1.3::3xflag* strain for 4 hours prior to our spermatogenesis time point. In the presence of SNPC-4 expression, SNPC-1.3 was enriched at both piRNA clusters, albeit to a lesser degree at the small cluster, and this enrichment was lost upon SNPC-4 depletion (Figures 3A and S4A). These data indicate that SNPC-1.3 binds piRNA loci during spermatogenesis in a SNPC-4-dependent manner *in vivo*.

**Figure 3.**
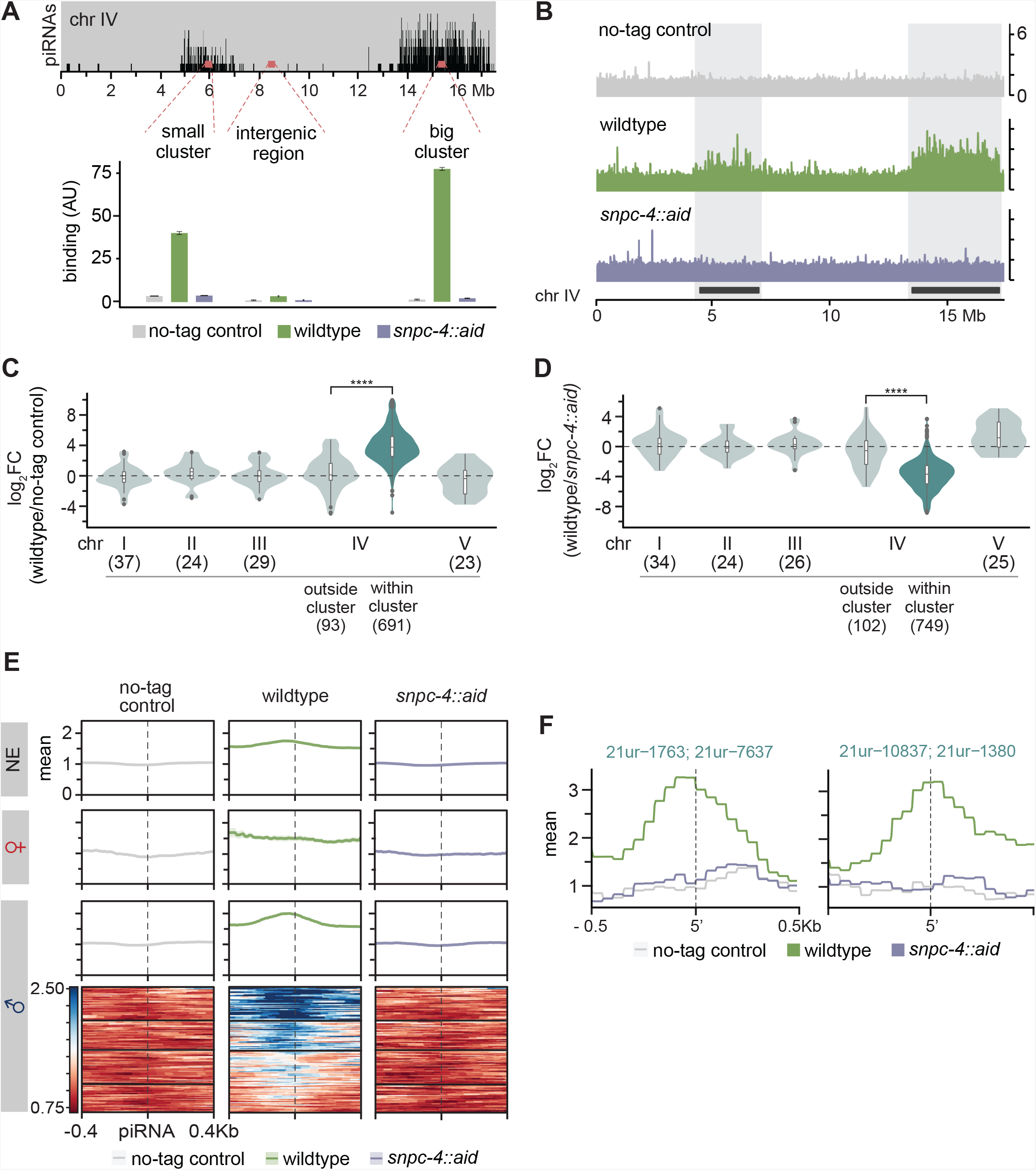
SNPC-1.3 binds male piRNA loci in a SNPC-4-dependent manner. (A) SNPC-1.3 binding at the piRNA clusters requires SNPC-4. SNPC-1.3::3xFlag binding normalized to input (mean ± SD of two technical replicates) on chromosome IV by ChIP-qPCR in a no-tag control, the strain expressing SNPC-4::3xFlag (wildtype), and in the strain expressing SNPC-4::3xFlag::AID, which undergoes TIR-1-mediated degradation by addition of auxin (*snpc-4::aid*). These labels (no-tag, wild-type, and *snpc-4::aid*) apply throughout Figure 3. Top panel depicts the density of piRNAs on chromosome IV with piRNAs predominantly found in the small (4.5–7 Mb) and big (13.5–17.2 Mb) cluster. (B) SNPC-1.3 binding profiles across chromosome IV in no-tag control, wildtype, and *snpc-4::aid*. The locations of the two piRNAs clusters are highlighted. (C) SNPC-1.3 binding is enriched at piRNA clusters on chromosome IV. SNPC-1.3-bound regions are enriched within piRNA clusters compared to regions outside of the piRNA clusters on chromosome IV (**** p ≤ 0.0001, Wilcoxon rank sum test). The number of bins analyzed is listed in parentheses. (D) SNPC-1.3 enrichment at piRNA clusters is dependent on SNPC-4. SNPC-1.3-bound regions within piRNA clusters are depleted compared to regions outside of the piRNA clusters on chromosome IV upon loss of SNPC-4 (****p ≤ 0.0001, Wilcoxon rank sum test). The number of bins analyzed is listed in parentheses. (E) Distribution of SNPC-1.3 reads (mean density ± standard error) around the 5’ nucleotide of mature piRNAs at the piRNA clusters. To resolve SNPC-1.3 binding between male and female piRNAs despite the high density of piRNAs, we selected 1 kb bins with all male (100), female (19), or non-germline enriched (279) piRNAs. (F) Examples of SNPC-1.3 binding at two regions containing two male piRNA loci. Regions are anchored on the 5’ nucleotide of each mature male piRNA and show mean read density ± standard error.

To examine the genome-wide binding profile of SNPC-1.3 and its dependency on SNPC-4, we performed ChIP-seq of wildtype, *snpc-1.3::3xflag*, and *snpc-1.3::3xflag; snpc-4::aid; P_sun-1_::TIR1* worms during spermatogenesis (Figures S4B–C). Consistent with our ChIP-qPCR results, we found that SNPC-1.3 binds piRNA clusters in a SNPC-4-dependent manner (Figures 3B and S4D). By quantifying the SNPC-1.3 signal over consecutive, non-overlapping 1 kb bins across the entire genome, we identified 691 1 kb regions within the chromosome IV piRNA clusters that were enriched for SNPC-1.3 in *snpc-1.3::3xflag* worms (wild-type) compared to the no-tag control worms (Figures 3C and S4E). Relative to *snpc-1.3::3xflag* (wild-type), worms depleted of SNPC-4 showed loss of SNPC-1.3 in 749 1 kb regions on chromosome IV piRNA clusters (Figure 3D). Furthermore, SNPC-1.3 enrichment (p< 2.2 × 10^−16^) and depletion (p< 2.2 × 10^−16^) were specific to the piRNA clusters on chromosome IV, and more than half (393/691) of the SNPC-1.3-enriched regions in *snpc-1.3::3xflag* worms were depleted upon degradation of SNPC-4 (Figures 3C–D, S4E–F).

To determine whether SNPC-1.3 preferentially binds male piRNA loci, we characterized the SNPC-1.3 signal around individual 5’ nucleotides of mature piRNAs. Again, we classified piRNAs as enriched in male or female germlines based on our small RNA-seq analysis in wild-type worms during spermatogenesis and oogenesis (Figure 2C). We found that SNPC-1.3 binding at male piRNA loci was most enriched just upstream of the piRNA 5’ nucleotide, which overlaps the conserved core motif (Figures 3E and S4G). This binding profile was very distinct for 1 kb bins that contained only male piRNAs (Figures 3F and S4H). Upon depletion of SNPC-4, this peak in male piRNAs was lost (Figures 3E and S4G). Although the binding profiles for individual female piRNAs exhibited more variability, there was little evidence for SNPC-1.3 binding and dependency on SNPC-4 at female loci (Figures 3E and S4G). Compared to the binding profile in male piRNA loci, SNPC-1.3 binding was observed to a lesser extent in piRNAs not enriched in the male and female germline (Figures 3E and S4G). Taken together, these observations indicate that SNPC-1.3 requires the core piRNA factor SNPC-4 to bind predominantly at male piRNA promoters.

### TRA-1 represses *snpc-1.3* and male piRNAs expression during oogenesis

As male piRNA expression and SNPC-1.3 protein expression are largely restricted to the male germline, we asked how SNPC-1.3 expression is regulated across development. *C. elegans* hermaphrodites produce sperm during the L4 stage and transition to producing oocytes as adults. To understand the mRNA expression profile of *snpc-1.3* relative to *snpc-4* and other developmentally regulated genes, we performed qRT-PCR across hermaphrodite development. *snpc-4* expression peaked during young adult and adult stages when oogenesis occurs (Figures 4A and S1E). These data suggest that low levels of SNPC-4 are sufficient for activating male piRNA biogenesis during spermatogenesis (Figure 1A). Consistent with SNPC-1.3 protein expression (Figure 1D), we observed specific *snpc-1.3* mRNA enrichment from L3 to early L4 stages, during spermatogenesis (Figure 4A). Given that *snpc-1.3* expression across development is regulated at the mRNA level, we examined the sequences upstream of the *snpc-1.3* coding region to identify potential *cis*-regulatory motifs. Less than 200 bp upstream of the *snpc-1.3* start codon, we identified three consensus binding sites for TRA-1 (Figure 4B), a transcription factor that controls the transition from spermatogenesis to oogenesis (Berkseth et al., 2013; Clarke and Berg, 1998; Zarkower and Hodgkin, 1993).

**Figure 4.**
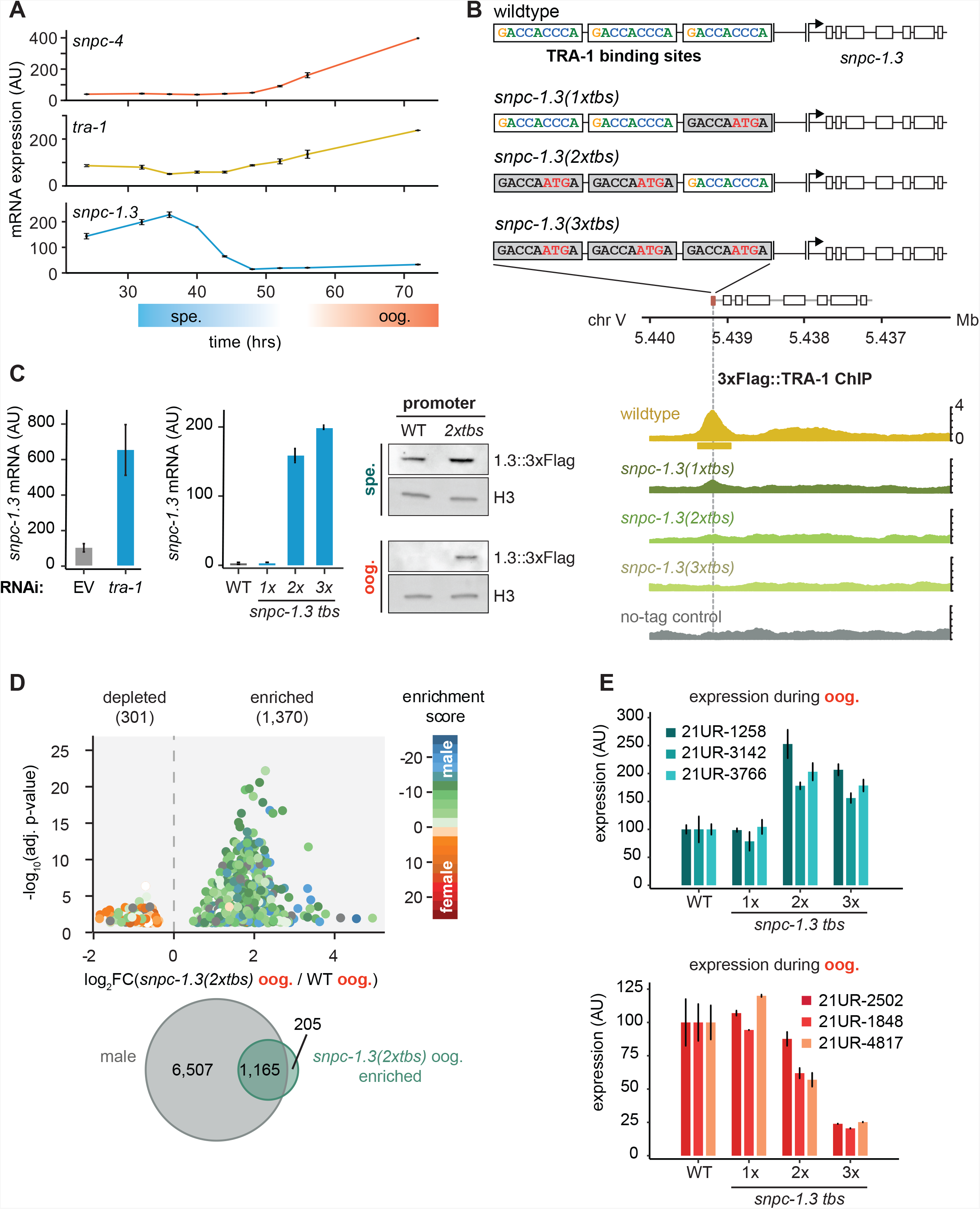
TRA-1 represses *snpc-1.3* and male piRNAs expression during oogenesis. (A) *snpc-1.3* mRNA levels peak during early spermatogenesis (spe.) while *tra-1* mRNA levels are highest during oogenesis (oog.). qRT-PCR and quantification of *snpc-1.3, snpc-4,* and *tra-1* mRNA normalized to *eft-2* mRNA across hermaphrodite development. Error bars: ± SD of two technical replicates. (B) TRA-1 binds to the *snpc-1.3* promoter. Schematic of the three TRA-1 binding sites upstream of the *snpc-1.3* locus in wildtype (top). Site-specific mutations shown in red were made in one, two, or three of the TRA-1 binding sites (grey denotes the mutated motifs). (Bottom) TRA-1 binding is reduced in TRA-1 binding site mutants assayed by TRA-1 ChIP-seq. (C) TRA-1 represses *snpc-1.3* mRNA expression during oogenesis. (Left) *snpc-1.3* mRNA expression is drastically upregulated upon RNAi-mediated knockdown of *tra-1* and (middle) in strains bearing mutations in two (*2xtbs*) or three (*3xtbs*) TRA-1 binding sites. Error bars indicate ± SD from two technical replicates. (Right) Western blot of SNCP-1.3::3xFlag expression driven under the wild-type and *2xtbs* mutant promoter during spermatogenesis (spe.) (top) and oogenesis (oog.) (bottom). H3 was used as the loading control. (D) A subset of male piRNAs are ectopically expressed during oogenesis in *snpc-1.3(2xtbs)* mutants. (Top) Volcano plot showing differential piRNA expression between *snpc-1.3(2xtbs)* mutants versus wildtype during oogenesis (oog.). piRNAs are colored by enrichment scores from Billi et al., 2013. (Bottom) Overlap of male piRNAs defined in Figure 2C with upregulated piRNAs in *snpc-1.3(2xtbs)* mutants. (E) Mutations at two (*2xtbs*) or three (*3xtbs*) TRA-1 binding sites enhance male piRNA expression (top) but attenuate female piRNA expression (bottom) during oogenesis. Error bars indicate ± SD from two technical replicates.

In the germline, TRA-1, a Gli family zinc-finger transcription factor, controls the sperm-to-oocyte decision by repressing both *fog-1* and *fog-3,* which are required for controlling sexual cell fate (Berkseth et al., 2013; Chen and Ellis, 2000; Lamont and Kimble, 2007; Zarkower and Hodgkin, 1993). Loss-of-function *tra-1* hermaphrodites exhibit masculinization of the female germline and develop phenotypically male-like traits (Hodgkin, 1987). We used RNAi to knock down *tra-1* and observed significant ectopic upregulation of *snpc-1.3* mRNA during oogenesis (Figure 4C). However, this upregulation of *snpc-1.3* expression could be an indirect effect of masculinization of the germline. Therefore, to test whether TRA-1 directly regulates *snpc-1.3*, we generated strains harboring mutations at the three TRA-1 binding sites (*tbs*) in the endogenous *snpc-1.3* promoter. Specifically, we mutated one (*1xtbs*), two (*2xtbs*), or all three (*3xtbs*) consensus TRA-1 binding motifs (Figure 4B). Disruption of the TRA-1 binding sites led to reduced TRA-1::3xFlag binding upstream of *snpc-1.3* as revealed by ChIP-seq, with the *3xtbs* mutant showing the greatest reduction of binding (Figures 4B and S5D). In addition, *snpc-1.3* mRNA levels were highly upregulated when multiple TRA-1 binding sites were mutagenized (Figure 4C), consistent with TRA-1 directly repressing *snpc-1.3* transcription during oogenesis. To confirm that SNPC-1.3 protein expression was also elevated in TRA-1 binding site mutants, we used CRISPR/Cas9 engineering to add a C-terminal 3xFlag tag at the *snpc-1.3* locus in *snpc-1.3(2xtbs)* mutants. Indeed, SNPC-1.3::3xFlag showed increased expression in the *snpc-1.3::3xFlag(2xtbs)* mutant during spermatogenesis and especially oogenesis (Figure 4C). Taken together, these findings show that TRA-1 binds to the *snpc-1.3* promoter to repress its transcription during oogenesis.

Given that *snpc-1.3* is robustly de-repressed during oogenesis in TRA-1 binding site mutants, we hypothesized that male piRNAs would also be ectopically upregulated during oogenesis. To test this, we performed small RNA-seq and compared piRNA levels in wildtype and *snpc-1.3(2xtbs)* worms during oogenesis (Table S3, Figure S5B–C, S5E). Using a FDR of ≤ 0.05, we saw significant upregulation of 1,370 piRNAs in *snpc-1.3(2xtbs)* mutants when compared to wildtype (Figure 4D). The majority of these upregulated piRNAs overlap with the male piRNAs that we identified in wildtype worms (Figure 4D). We also confirmed this result by Taqman qPCR analysis, which showed that male piRNAs were significantly upregulated in *snpc-1.3(2xtbs)* and *snpc-1.3(3xtbs)* mutants compared to wildtype during oogenesis (Figure 4E). Taken together, these data suggest that TRA-1 directly binds to *tbs* sites in the *snpc-1.3* promoter to repress its transcription and consequently, male piRNA expression during oogenesis.

Our data showed that female piRNAs are upregulated during spermatogenesis upon loss of *snpc-1.3* (Figure 2D). Consistent with this result, we found that female piRNAs show reduced expression during oogenesis upon upregulation of SNPC-1.3 in *snpc-1.3(2xtbs)* and *snpc-1.3(3xtbs)* mutants compared to wildtype (Figure 4E). We posit that SNPC-1.3 plays a direct role in activating male piRNA transcription, while indirectly limiting female piRNA transcription by sequestering core piRNA transcription factors to male piRNA loci.

### SNPC-1.3 is critical for male fertility

Given the global depletion of male piRNAs in *snpc-1.3(-)* mutants and the progressive fertility defects seen in *prg-1(-)* mutants (Batista et al., 2008; Wang and Reinke, 2008), we hypothesized that *snpc-1.3(-)* worms might also show fertility defects. Indeed, we found that *snpc-1.3(-)* hermaphrodites exhibited significantly reduced fertility compared to wildtype when grown at 25°C (Figure 5A). To address whether this decreased fertility was due to defects during spermatogenesis or oogenesis, we compared brood sizes from crosses of *fem-1(-)* females and *him-8(-)* males with or without *snpc-1.3*. Compared to *him-8(-)* males, we found that *him-8(-); snpc-1.3(-)* males generated significantly smaller brood sizes when crossed with *fem-1(-)* females; in contrast, *fem-1(-); snpc-1.3(-)* and *fem-1(-)* females generated similar brood sizes when crossed with *him-8(-)* males (Figure 5B). As an orthogonal test, we crossed hermaphrodites with transgenic males that express the COL-19::GFP marker in the cuticle to facilitate counting of cross progeny (Figure S6A). All *col-19::gfp; snpc-1.3(-)* males produced fewer GFP+ progeny than wild-type *col-19::gfp* males, whereas wildtype or *snpc-1.3(-)* hermaphrodites generated similar numbers of GFP+ progeny when crossed with wild-type *col-19::gfp* males (Figure S6A). These results suggest that the reduced fertility of *snpc-1.3(-)* mutants likely reflect defects during spermatogenesis.

**Figure 5.**
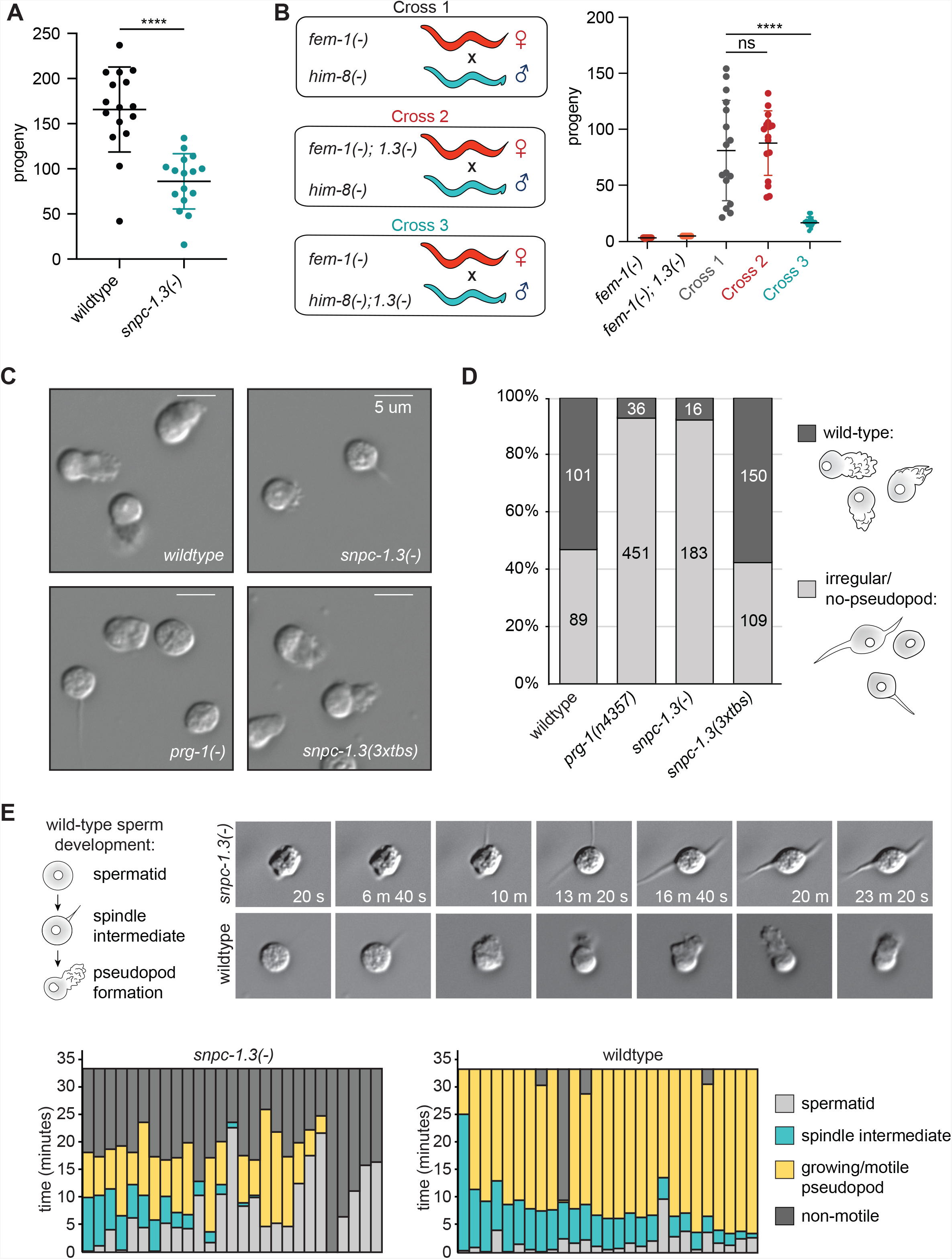
SNPC-1.3 is critical for male fertility. (A) *snpc-1.3(-)* hermaphrodites exhibit sterility at 25°C. Circles correspond to the number of viable progeny from singled hermaphrodites (n=16). Black bars indicate mean ± SD. Statistical significance was assessed using Welch’s t-test (****p ≤ 0.0001) (B) *snpc-1.3* promotes male fertility but is dispensable for female fertility (Left) Diagram illustrates crosses between strains for mating assays (*1.3(-)* denote*s snpc-1.3(-)*). (Right) *snpc-1.3(-)*; *him-8(-)* males crossed to *fem-1*(-) females show severe fertility defects (Cross 3). *snpc-1.3; fem-1(-)* females crossed to *him-8(-)* males (Cross 2) show equivalent fertility similar to *fem-1(-)* females crossed to *him-8(-)* males (Cross 1). Circles correspond to the number of viable progeny from cross (n=16). Black bars indicate mean ± SD. Statistical significance was assessed using Welch’s t-test (ns: not significant; ****p ≤ 0.0001) (C) *snpc-1.3(-)* spermatids exhibit severe morphological defects. Images of pronase-treated sperm of wildtype, *prg-1(-), snpc-1.3(-)*, and *snpc-1.3(2xtbs)* males. (D) *snpc-1.3(-)* spermatids exhibit severe sperm maturation defects. (E) (Top) Images depicted at 3 min intervals of a sperm undergoing activation and maturation. Imaging of spermatid commenced ~3 min after pronase treatment. (Bottom) Graphical display of individual sperm tracked over time after pronase treatment.

To investigate the cause of *snpc-1.3*-dependent loss of male fertility, we examined spermiogenesis and sperm morphology in *snpc-1.3(-)* males in greater detail. After meiotic differentiation in the male germline, male spermatids are induced by ejaculation and undergo spermiogenesis, a process that converts immature spermatids to motile sperm with a functioning pseudopod. Spermiogenesis can be induced *in vitro* by isolating spermatids from the spermatheca and treating them with pronase (Shakes and Ward, 1989). Males lacking *prg-1* still generate differentiated spermatids, but rarely produce normal pseudopodia upon activation (Figures 5C and 5D) (Wang and Reinke, 2008). Similar to *prg-1(-)* mutants, *snpc-1.3(-)* spermatids were rarely able to form normal pseudopodia. In contrast, *snpc-1.3(3xtbs)* sperm formed normal pseudopodia at a frequency similar to wildtype (Figures 5C–D). In addition, many of the *snpc-1.3(-)* spermatids resembled sperm undergoing intermediate stages of spermiogenesis. Spermiogenesis *in vivo* starts off with spherical spermatids that enter into an intermediate stage characterized by the growth of spiky protrusions. This stage is then followed by fusion of the spiky protrusions into a motile pseudopod (Figure 5E). To understand the dynamics of *snpc-1.3(-)* sperm progression through spermiogenesis, we treated spermatids with pronase and observed each activated spermatid over time. Wild-type spermatids spent an average of 6.2 min ± 4.5 min in the intermediate state before polarization and pseudopod development. In contrast, *snpc-1.3(-)* spermatids occupied the intermediate state for a significantly shorter period of time (2.9 min ± 3.7 min, p<0.05; Student’s t test) before forming pseudopods. By tracking each individual spermatid across spermiogenesis, we found most *snpc-1.3(-)* spermatids were unable to sustain motility. While wild-type spermatids exhibited motility for an average of 24 min ± 10.35 min, *snpc-1.3(-)* spermatids showed motility for significantly shorter period of time (7.3 min ± 5.7 min, p<0.05; Student’s t test) before becoming immotile (Figure 5E, bottom). These results indicate that *spnc-1.3(-)* males have defective spermatogenesis processes and exhibit similar fertility defects as *prg-1(-)* mutants.

## DISCUSSION

Based on our data, we propose that *C. elegans* SNPC-1.3, a human SNAPC1 ortholog, is a male piRNA transcription factor. SNPC-1.3 preferentially binds to the promoters of male piRNAs (Figure 3) and is critical for their expression (Figure 2). SNPC-1.3 expression, reflecting the developmental profile of male piRNAs (Figure S5A), is highest during spermatogenesis. We demonstrate that the *snpc-1.3* locus itself is regulated by the sex determination regulator, TRA-1 (Figure 4). During spermatogenesis, *tra-1* expression is low, and *snpc-1.3* and other male-promoting genes are licensed for expression. In contrast, *tra-1* expression is upregulated during oogenesis and TRA-1 binds the *snpc-1.3* promoter to repress its transcription, promoting the expression of female over male piRNAs (Figure 6). We propose that SNPC-1.3, via its interaction with SNPC-4, can switch the specificity of the core piRNA complex from binding at female to male piRNA loci.

**Figure 6.**
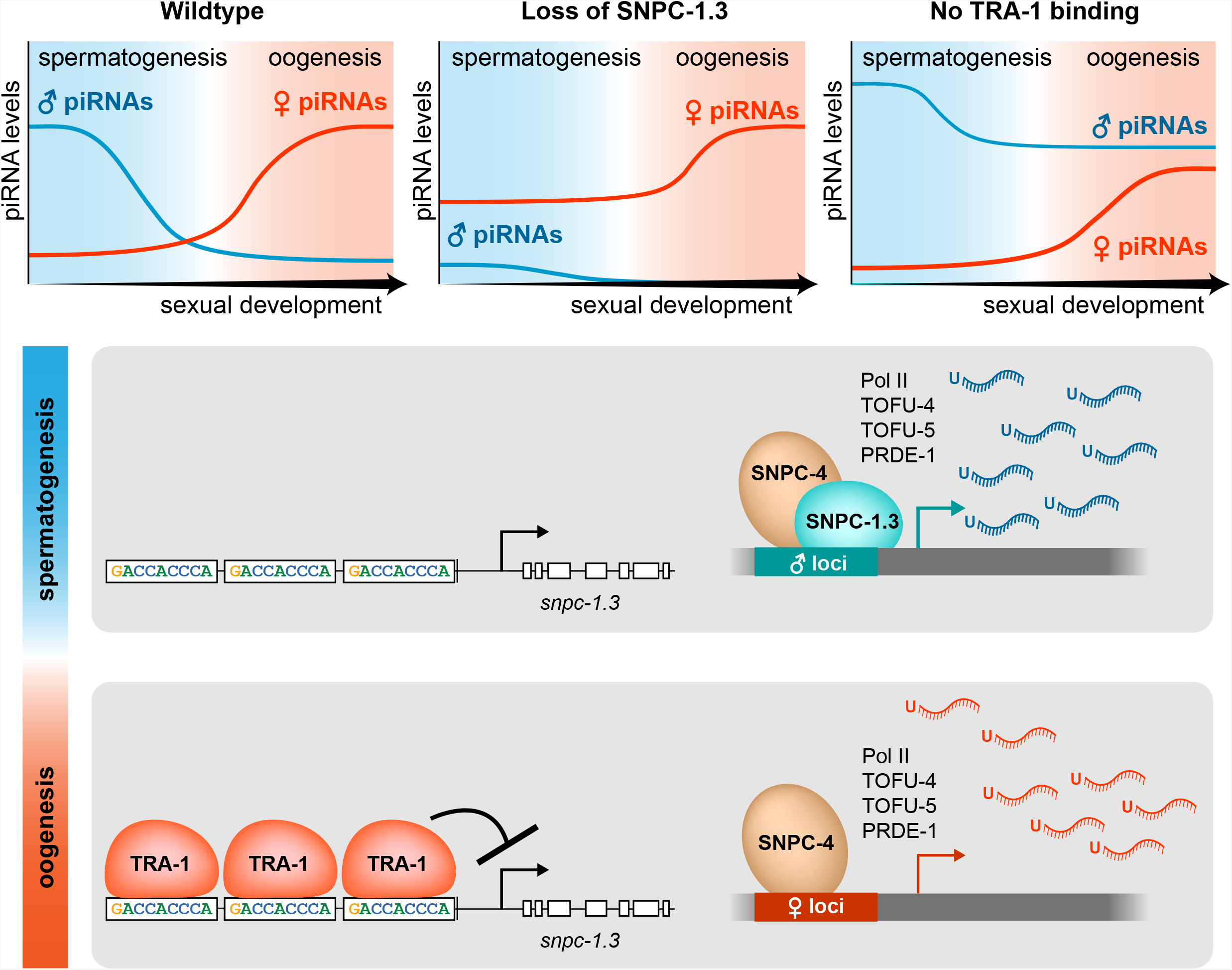
Model illustrating the dynamics of male and female piRNA transcription across *C. elegans* sexual development. In wild-type worms, male piRNA and female piRNA expression peaks during spermatogenesis and oogenesis, respectively. (Top) In *snpc-1.3(-)* mutants, male piRNA expression is abrogated, and female piRNA expression is moderately enhanced across sexual development relative to wildtype. In TRA-1 binding site mutants, *snpc-1.3* expression is de-repressed causing ectopic upregulation of male piRNAs and moderate repression of female piRNA expression during oogenesis relative to wildtype. (Bottom) During spermatogenesis, SNPC-1.3 interacts with SNPC-4 at male piRNA promoters regions to drive male piRNA transcription. During oogenesis, TRA-1 represses the transcription of *snpc-1.3* which results in the suppression of male piRNA transcription, thus leading to enhanced transcription of female piRNAs.

### How is the expression of male and female piRNAs coordinated?

Given its role as a putative male piRNA transcription factor, we expected that deletion of *snpc-1.3* would result in loss of male piRNAs but with no consequences to the expression of female piRNAs. However, loss of *snpc-1.3* also results in increased female piRNA expression during spermatogenesis (Figure 2), whereas ectopic overexpression of *snpc-1.3* during oogenesis leads to decreased female piRNA levels (Figure 4). Taken together, our findings suggest that male and female piRNA transcription are not completely separable from each other and that the balance in expression of the two piRNA subclasses may be dictated by the allocation of shared core transcription factors such as SNPC-4.

Similar to multiple gene classes activated by general transcription factors (Levine et al., 2014), we speculate that male and female piRNA promoters compete for access to a limited pool of the core biogenesis complex, which includes SNPC-4, PRDE-1, TOFU-4, and TOFU-5 (Figure 1). Therefore, we propose a model in which the expression and binding of SNPC-1.3 to core piRNA factors serves to “sequester” the core complex away from female promoters. Mechanistically, we posit that the core piRNA transcription complex is specified to female promoters, and that only upon association with SNPC-1.3 is the core machinery directed to male piRNA promoters. We predict that when SNPC-1.3 is absent, more SNPC-4 and other previously identified cofactors are available to transcribe female piRNAs. Conversely, overexpression of SNPC-1.3 leads to the disproportionate recruitment of the core machinery to male promoters, leading to the indirect downregulation of female piRNAs. By controlling male piRNA expression, SNPC-1.3 is crucial for maintaining the balance between male and female piRNA levels across development.

While the default specification of the core complex to female promoters presents perhaps the most parsimonious explanation underlying male and female piRNA expression, we cannot exclude the possibility that an additional female-specific *trans*-acting factor may direct the core piRNA complex to female promoters. If true, we speculate that the developmental expression of such a factor (low during spermatogenesis and high during oogenesis), coupled with the developmental expression of SNPC-1.3, would coordinate the differential expression of male and female piRNAs. During spermatogenesis, SNPC-1.3 is more highly expressed such that the core machinery would primarily be directed to male promoters. In contrast, during oogenesis, SNPC-1.3 expression is low, concomitant with elevated expression of a female factor to license transcription of female piRNAs. This model, where both factors are present during both spermatogenesis and oogenesis but in different ratios, would also be consistent with our piRNA expression analysis in *snpc-1.3* loss-of-function and overexpression mutants.

### The piRNA pathway co-opts snRNA biogenesis machinery

Our work adds to a growing body of evidence that snRNA machinery has been hijacked at multiple stages in *C. elegans* piRNA biogenesis, including transcription (Kasper et al., 2014; Weng et al., 2018) and termination (Beltran et al., 2019). Investigating potential parallels between snRNA and piRNA biogenesis may provide useful clues into the role of SNPC-1.3 in the piRNA complex.

The minimal snRNA SNAPc complex consists of a 1:1:1 heterotrimer of the subunits SNAPC4, SNAPC1, and SNAPC3 in humans and SNAP190, SNAP43, and SNAP50 in flies (Henry et al., 1998; Hung and Stumph, 2010; Li et al., 2004; Ma and Hernandez, 2002; Mittal et al., 1999) (Figure 1B). *In vitro* studies have shown that the trimer must assemble before the complex is able to bind DNA. Similarly, our data show SNPC-1.3 requires SNPC-4 to bind at the piRNA clusters (Figure 3). We speculate the piRNA complex is assembled in a similar fashion to the snRNA complex. Based on this model, we expect that SNPC-4 binding at male piRNA loci is abolished in a *snpc-1.3* mutant. However, conclusive evidence that SNPC-4 binding at male piRNA promoters requires SNPC-1.3 is still lacking. Due to the highly clustered nature of *C. elegans* piRNAs, we have been unable to discriminate detectable differences in SNPC-4 binding between male and female piRNAs in *snpc-1.3(-)* mutants, as assayed by traditional ChIP-seq methods. Application of higher resolution techniques may be required to address this question.

Given that piRNAs have co-opted *trans*-acting factors from snRNA biogenesis (Kasper et al., 2014), it would not be surprising if piRNAs also co-evolved *cis*-regulatory elements for transcription factor binding from snRNA loci. Recently, Beltran et al. (2019) identified similarity between the 3’ end of PSEs of snRNA promoters and the 8 nt piRNA core motif in nematodes. In addition, Pol II and Pol III transcription from snRNA promoters share a common PSE, but are distinguished by the presence of other unique motifs (Hung and Stumph, 2010). Correspondingly, the canonical Type I and less abundant Type II piRNAs can be discriminated by the presence or absence of the 8 nt core motif, respectively. Factors such as TOFU-4 and TOFU-5 function in both Type I and II piRNA expression, whereas PRDE-1 is only required for Type I piRNAs (Kasper et al., 2014; Weng et al., 2018). Altogether, these observations highlight the importance of *cis*-regulatory elements in specifying the expression of snRNAs and piRNA classes. In addition to enrichment of cytosine at the 5’ position in the male core motif (Billi et al., 2013), we hypothesize that as-yet unidentified motifs may further discriminate male from female piRNA promoters. While we observed SNPC-1.3 binding to be enriched upstream of male piRNA loci (Figure 3), we cannot definitively conclude that SNPC-1.3 binds to the male-specific core motif, given the limitations of conventional ChIP-seq in resolving the SNPC-1.3 footprint. Identifying the factors that specifically bind the 8 nt core motif and other potential *cis*-regulatory elements important for sex-biased piRNA expression will require further investigation.

### What are the functions of male piRNAs in *C. elegans*?

Our data suggest that SNPC-1.3 is essential for proper spermiogenesis (Figure 5). We hypothesize the global loss of male piRNAs in a *snpc-1.3(-)* mutant is responsible for the higher incidence of spermiogenesis arrest and subsequent loss in fertility, although it is possible that SNPC-1.3 may have other or additional effects on male fertility. Characterization of *prg-1(-)* mutants during spermiogenesis agree with our findings that loss of piRNAs in the male germline leads to acute defects directly responsible for fertility (Wang and Reinke, 2008). Since the initial discovery of piRNA function in the targeting and silencing of transposons in *Drosophila* (Vagin et al., 2006; Brennecke et al., 2007), analyses in other systems have revealed that piRNAs have acquired neofunctions at later points along the evolutionary timescale (Ozata et al., 2019).

While it is estimated that as much as 45% of the human genome encodes for transposable elements (McCullers and Steiniger, 2007), only 12% of *C. elegans* genome encodes such elements. Furthermore, nearly all of these regions are inactive in *C. elegans* (Bessereau, 2006). In contrast to *Drosophila* piRNAs that target and silence transposons with perfect complementarity (Brennecke et al., 2007), *C. elegans* piRNAs are thought to bind a broad range of endogenously expressed transcripts by partial complementarity (Ashe et al., 2012; Shen et al., 2018; Zhang et al., 2018). Together, these findings suggest that worm piRNAs function in capacities distinct from canonical transposon silencing. While a recent methodology used cross-linking, ligation, and sequencing of piRNA:target hybrids (CLASH) to determine that female piRNAs engage with almost every germline transcript (Shen et al., 2018), how male piRNAs select their targets has yet to be examined. Like piRNAs characterized in the female germline, male piRNAs may be interfacing with a broad range of targets to regulate gene expression for proper spermatogenesis. Loss of *prg-1* in males causes the downregulation of a subset of spermatogenesis-specific genes (Wang and Reinke, 2008), suggesting male piRNAs serve a protective function for spermatogenic processes. The characterization of the *in vivo* landscape of male piRNA target selection using CLASH may provide insights into piRNA function during spermatogenesis.

### Why are male piRNAs restricted from the female germline by TRA-1?

Sperm and oocytes pass epigenetic information such as noncoding RNAs to the next generation (Hammoud et al., 2014; Brykczynska et al., 2010; Tabuchi et al., 2018; Kaneshiro et al., 2019). Recent studies show maternal piRNAs trigger the production of endo-siRNAs, called 22G RNAs for their 5’ bias for guanine and 22 nt length, to transmit an epigenetic memory of foreign versus endogenous elements to the next generation (Ashe et al., 2012; Buckley et al., 2012; Shirayama et al., 2012). We predict that misexpression of male piRNAs in the female germline may perturb the native pool of female piRNAs necessary for appropriate recognition of self versus non-self elements. This may explain the decrease in fertility we observed in multiple TRA-1 binding site mutant hermaphrodites (Figure S6B). As *snpc-1.3(3xtbs)* sperm do not seem to exhibit significant morphological defects (Figure 5), the fertility defects in the *snpc-1.3(3xtbs)* mutants could be due to problems arising in oogenesis. However, based on our sequencing data in *snpc-1.3(2xtbs)* mutants, we cannot distinguish whether fertility defects during oogenesis are due to upregulation of male piRNAs, downregulation of female piRNAs, a combination of the two, or misexpression of downstream endo-siRNAs triggered by piRNAs. Further study of *snpc-1.3* gain-of-function mutants in oogenesis will enhance our understanding of the physiological consequences of expressing male piRNAs in the female germline.

### The intersection between sex determination and sex-specification of piRNA expression

We speculate that gene duplication of the *snpc-1* family of genes occurred early during nematode evolution and allowed for the acquisition of new functions by *snpc-1* paralogs, specifically, from snRNA to piRNA biogenesis. At least two SNPC-1 paralogs are present within the distantly related nematode species, *Plectus sambesii.* Furthermore, we predict that the co-option of SNPC-1 paralogs for piRNA biogenesis may have occurred in parallel with the evolution of the nematode sex determination pathway. TRA-1 is a sex determination factor that acts to repress male-promoting gene expression in female germ cells to promote female germ cell fate. While *Drosophila* sex determination utilizes different factors than *C. elegans*, further investigation into the conservation of TRA-1 shows that it is a common feature in at least the nematode lineage (Pires-daSilva and Sommer, 2004). Additionally, just as we have shown that TRA-1 represses *snpc-1.3* in *C. elegans* (Figure 4), TRA-1 binding motifs GGG(A/T)GG are present in the putative upstream promoter regions of *snpc-1.3* homologs identified in *C. briggsae, C. brenneri* and *C. nigoni* (Figure S6C). Taken together, these analyses point to a conserved link between sex determination and piRNA biogenesis pathways among nematodes.

In summary, our work reveals that SNPC-1.3 is specified to the male germline and is essential for male piRNA expression. We have identified SNPC-1.3 as a major target of TRA-1 repression in the female germline. Future studies will likely uncover additional factors required to coordinate the proper balance of sex-specific piRNAs required for proper germline development and animal fertility.

## CONTACT FOR REAGENT AND RESOURCE SHARING

Further information and requests for resources and reagents should be directed to and will be fulfilled by the Lead Contact, John K. Kim.

## EXPERIMENTAL MODEL AND SUBJECT DETAILS

*C. elegans* strains were maintained at 20°C according to standard procedures (Brenner, 1974), unless otherwise stated. Bristol N2 was used as the wildtype strain. Except for RNAi and ChIP experiments, worms were fed *E. coli* strain OP50. Worms used for ChIP were fed *E. coli* strain HB101.

## METHOD DETAILS

### Generations of strains

CRISPR/Cas9-generated strains were created as described in Paix et al., 2015 and are listed in Table S4. crRNA and repair template sequences of CRISPR generated strains are listed in Table S4. After initial phenotyping of *snpc-1.3a::3xflag* and *snpc-1.3b::3xflag* (Figure S1), *snpc-1.3a::3xflag* was used for all subsequent experiments (and is referred to as *snpc-1.3::3xflag*).

### RNAi assays

Bacterial RNAi clones were grown from the Ahringer RNAi library (Kamath and Ahringer, 2003). All clones used are listed in the Table S4. Synchronized L1 worms were plated on HT115 bacteria expressing dsRNA targeting the gene interest or L4440 empty vector as a negative control as previously described in (Timmons and Fire, 1998). All RNAi experiments were performed at 20°C unless otherwise stated.

### RNA extraction, library preparation, and sequencing

After hypochlorite preparation and hatching in M9 buffer, *snpc-4::aid::ollas* and *snpc-4::aid::ollas; P_sun-1_::TIR1* worms were transferred from NGM plates to plates containing 250 μM auxin 20 h before collection of L4 and gravid worms, 48 and 72 h after plating L1 worms, respectively. Worms were collected in TriReagent (Thermo Fisher Scientific) and subjected to three freeze-thaw cycles. Following addition of 1-bromo-3-chloropropane (BCP), the aqueous phase was then precipitated with isopropanol at −80°C for 2 h. To pellet RNA, samples were spun at 21,000 x *g* for 30 min at 4°C. After three washes in 75% ethanol, the pellet was resuspended in water.

RNA concentration and quality was measured using a TapeStation (Agilent Technologies). 16–30 nt small RNAs were size-selected from 5 μg total RNA on 17% denaturing polyacrylamide gels. Small RNAs were treated with 5’ polyphosphatase (Illumina) to reduce 5’ triphosphate groups to monophosphates to enable 5’ adapter ligation. Small RNA-sequencing libraries were prepared using the NEBNext® Multiplex Small RNA Library Prep Set for Illumina (NEB). Small RNA amplicons were size-selected on 10% polyacrylamide gels and quantified using qRT-PCR. Samples for each developmental time point were pooled into a single flow cell and single-end, 75 nt reads were generated on a NextSeq 500 (Illumina). An average of 42.01 million reads (range 33.05–50.39 million) was obtained for each library.

### Quantitative RT-PCR

Taqman cDNA synthesis was performed as previously described (Weiser et al., 2017). Briefly, for quantification of piRNA levels, TaqMan small RNA probes were designed and synthesized by Applied Biosystems. All piRNA species assessed by qPCR were normalized to U18 small nucleolar RNA. 50 ng of total RNA was used for cDNA synthesis. cDNA was synthesized by Multiscribe Reverse Transcriptase (Applied Biosystems) using the Eppendorf Mastercycler Pro S6325 (Eppendorf). Detection of small RNAs was performed using the TaqMan Universal PCR Master Mix and No AmpErase® UNG (Applied Biosystems). The sequences used for developing custom small RNA probes used for Taqman qPCR are listed in Table S4. For quantification of mRNA levels, cDNA was made using 500 ng of total RNA using Multiscribe Reverse Transcriptase (Applied Biosystems). Assays for mRNA levels were performed with Absolute Blue SYBR Green (ThermoFisher) and normalized to *eft-2* using CFX63 Real Time System Thermocyclers (Biorad). All qPCR primers used are listed in Table S4.

### Covalent crosslinking of Dynabeads

Protein G Dynabeads (ThermoFisher Scientific, 1003D) were coupled to monoclonal mouse anti-FLAG antibody M2 (Sigma Aldrich, F1804). After 3 washes in 1x PBST (0.1% Tween), Dynabeads were resuspended with 1x PBST with antibody, for a final concentration of 50 μg antibody per 100 μL beads. The antibody-bead mixture was nutated for 1 h at room temperature. After 3 washes in 1x PBST and 2 washes in 0.2 M sodium borate pH 9.0, beads were nutated in 22 mM DMP (Sigma Aldrich, D8388) in 0.2 M sodium borate for 30 min at room temperature. Following 2 washes in ethanolamine buffer (0.2 M ethanolamine, 0.2 M NaCl pH 8.5), beads were nutated for 1 h at room temperature in the same buffer. Beads were placed into the same volume of ethanolamine buffer as the starting bead volume for storage at 4°C until use.

### Immunoprecipitation for mass spectrometry and co-IP experiments

For SNPC-4 IP mass spectrometry, synchronized populations of ~200,000 *him-8(e1489)* L4s and ~50,000 *fem-1(hc17)* females were grown at 25°C and collected on OP50. For co-IP experiments, ~500,000 L4s and ~250,000 gravid worms were grown and collected from OP50 plates. After washing in M9, the gut was cleared for 15 min with nutation in M9 buffer before collection.

Unless otherwise stated, all samples for mass spectrometry and co-IP used in this study were subjected to the following procedure. After three washes in M9 and one wash in water, worms were frozen and ground using the Retsch MM400 ball mill homogenizer for 2 rounds of 1 min at 30s^−1^. Frozen worm powder was resuspended in 1x lysis buffer used previously (Moissiard et al., 2014) (50 mM Tris-HCl pH 8.0, 150 mM NaCl, 5 mM MgCl_2_, 1 mM EGTA, 0.1% NP-40, 10% glycerol) and protease inhibitor cocktail (Roche). After Bradford assay (ThermoFisher Scientific), lysates were normalized using lysis buffer and protease inhibitor. Benzonase (Sigma-Aldrich, E1014) was added to a final concentration of 1 μL/mL of lysate and nutated for 10 min at 4°C. After centrifugation for 10 min at 4,000 x *g*, 1 mL of supernatant was added to 50 μL of crosslinked Dynabeads and nutated for 15 min at 4°C. Samples were then washed 3 times in 1x lysis buffer with protease inhibitors before 1 h nutation in 50 μL of 2 mg/mL FLAG peptide (Sigma-Aldrich, F4799) diluted in 1x lysis buffer. Complete eluate, as well as 5% of crude lysate (after addition of benzonase), input, pellet, and post-IP samples were added to 2x Novex Tris-Glycine SDS Sample Buffer (ThermoFisher Scientific, LC2676) to 1x. Samples were then subjected to western blotting as described below.

### Mass spectrometry and analysis

Proteins were precipitated with 23% TCA and washed with acetone. Protein pellets solubilized in 8 M urea, 100 mM Tris pH 8.5, and reduced with 5 mM Tris(2-carboxyethyl)phosphine hydrochloride (Sigma-Aldrich, St. Louis, MO, product C4706) and alkylated with 55 mM 2-Chloroacetamide (Fluka Analytical, product 22790). Proteins were digested for 18 h at 37°C in 2 M urea 100 mM Tris pH 8.5, 1 mM CaCl_2_ with 2 μg trypsin (Promega, Madison, WI, product V5111). Single phase analysis (in replicate) was performed using a Dionex 3000 pump and a Thermo LTQ Orbitrap Velos using an in-house built electrospray stage (Wolters et al., 2001). Protein and peptide identification and protein quantitation were done with Integrated Proteomics Pipeline, IP2 (Integrated Proteomics Applications, Inc., San Diego, CA. http://www.integratedproteomics.com/). Tandem mass spectra were extracted from raw files using RawConverter (He et al., 2015) with monoisotopic peak option and were searched against protein database release WS260 from Wormbase, with FLAG-tagged SNPC-4, common contaminants and reversed sequences added, using ProLuCID (Peng et al., 2003; Xu et al., 2015). The search space included all fully-tryptic and half-tryptic peptide candidates with a fixed modification of 57.02146 on C. Peptide candidates were filtered using DTASelect (Tabb et al., 2002).

Using custom R scripts, average enrichment between SNPC-4::3xFlag and no-tag control immunoprecipitation experiments were calculated. For each experiment, enrichment was normalized by dividing the peptide count for each protein by the total peptide count. Adjusted *p-*values were calculated by applying the Bonferroni method using DESeq2 (Love et al., 2014).

### Western blotting

At least 250,000 L4 and 50,000 gravid worms were collected from OP50 plates. L4s were collected on empty vector L4440 or *snpc-1.3* RNAi. For *glp-4* temperature-shift experiments, worms grown at 15°C were egg prepped and hatched. Subsequent synchronized L1s were transferred to 25°C. After washing in M9, the gut was cleared for 15 min with nutation in M9. After three washes in M9 and one wash in water, worms were frozen and pulverized using the Retsch MM400 ball mill homogenizer for 2 rounds of 1 min at 30s^−1^. Frozen worm powder was resuspended in 1x lysis buffer used previously (Moissiard et al., 2014) (50 mM Tris-HCl pH 8.0, 150 mM NaCl, 5 mM MgCl_2_, 1 mM EGTA, 0.1% NP-40, 10% glycerol, and protease inhibitor cocktail (Roche)). After Bradford assay (ThermoFisher Scientific), lysates were normalized using lysis buffer. Benzonase (Sigma-Aldrich, E1014) was added to a final concentration of 1 μL/mL of lysate and nutated for 10 min at 4°C. The lysate was then added to 4x loading buffer (200 mM Tris pH 6.8, 8% SDS, 0.4% bromophenol blue, 40% glycerol, 20% beta-mercaptoethanol) to a final concentration of 1x. Samples were run on either 8–16% or 8% Novex WedgeWell Tris-Glycine precast gels (ThermoFisher), and transferred to PVDF membrane (Millipore). Mouse anti-Flag, rabbit anti-gamma tubulin, and rabbit anti-H3 were used at 1:1,000, 1:5,000, and 1:15,000, respectively. Anti-mouse and anti-rabbit (for tubulin) antibodies were used at 1:5,000. To blot for H3, anti-rabbit secondary was used at 1:15,000. Antibodies used were Sigma-Aldrich F1804 (mouse anti-Flag), Sigma-Aldrich T1450 (rabbit anti-gamma tubulin), and Abcam ab1791 (rabbit anti-H3), GE Healthcare NA931 (sheep anti-mouse), and Jackson Laboratories 111035045 (goat anti-rabbit). Both high sensitivity Amersham ECL Prime (GE Healthcare, RPN2232) (for SNPC-1.3 blotting) and regular sensitivity Pierce ECL (ThermoFisher, 32209) were used for exposure in a BioRad ChemiDoc Touch system.

### Chromatin immunoprecipitation, library prep, and sequencing

Worms were grown in liquid culture as previously described (Zanin et al., 2011). 250 μM auxin was added to *snpc-1.3::3xflag; snpc-4::aid::ollas; P_sun-1_::TIR1* worms 4 h before collection at 48 h post L1 at 20°C. After washing, the gut was cleared for 15 min by nutation in M9, followed by three washes in M9. Worms were live-crosslinked in 2.6% formaldehyde in water for 30 min at room temperature with nutation. Crosslinking was quenched with a final concentration of 125 mM glycine for 5 min with nutation. After three washes with water, worms were flash-frozen in liquid nitrogen. Frozen worm pellets were ground into powder using the Retsch MM40 ball mill homogenizer for 2 rounds of 1 min at 30s^−1^. Frozen worm powder was resuspended in 1x RIPA buffer (1xPBS, 1% NP-40, 0.5% sodium deoxycholate, 0.1% SDS) for 10 min at 4°C. Crosslinked chromatin was sonicated using a Diagenode Bioruptor Pico for three 3-min cycles, 30 sec on/off. 10 μg chromatin was nutated overnight at 4°C with 2 μg of Flag antibody (Sigma-Aldrich, F1804) and then for 1.5 h with 50 μL mouse IgG Dynabeads (Invitrogen). Input amount was 10% of IP. Chromatin was de-crosslinked and extracted as described previously (Weiser et al., 2017). Individual input and IP samples of each genotype were processed for both sequencing and quantitative PCR.

Libraries were prepared and multiplexed using the Ovation Ultralow Library Systems v2 (NuGEN Technologies) according to the manufacturer’s protocol. The Illumina HiSeq 4000 platform was used to generate 50 bp single-end reads for SNPC-1.3 ChIP-seq libraries. The NovaSeq 6000 platform was used to generate 50 bp paired-end reads for TRA-1 ChIP libraries.

### Quantitative PCR of ChIP samples

ChIP DNA was eluted in 18 μL of 1x TE pH 8.0 and 2 μL of 20 mg/mL RNase A (Invitrogen, Thermo Fisher Scientific). For a final reaction volume of 25 μL, each reaction consisted of final 1x Absolute Blue SYBR Green (Thermo Fisher Scientific), 35 nM each of forward and reverse primer, and 2 uL ChIP eluate. Reactions were performed in technical duplicates in a BioRad CF96 Real Time PCR thermal cycler.

### Hermaphrodite fertility assays

Gravid worms (previously maintained at 20°C) were subjected to hypochlorite treatment and their progeny were plated onto NGM at 25°C (P0). At the L2 or L3 stage, worms were singled onto individual plates and their progeny (F1) counted.

### Mating assays

To test male-dependent rescue of *fem-1(hc17)* fertility, 10–12 hermaphrodites of each strain were grown at 20°C and embryos were isolated by allowing egg lay for 2 h before removal. Embryos were shifted to 25°C and upon reaching the L4 stage (24 h), ten *him-8(e1489)* L4 males were transferred and mated with two *fem-1(hc17)* females. Brood size was quantified by counting when a majority of progeny had at least reached the young adult stage (about 3 days after transfer). To test the fertility of the hermaphrodites upon mating, 10–12 hermaphrodites of each strain were grown at 20°C and embryos were isolated after egg lay for 2 h before removal. Embryos were shifted to 25°C and ten *col-19(GFP+)* L4-staged males (24 h) were then transferred with a single hermaphrodite (36 h) and the number of live cross progeny were counted after reaching adulthood. Brood size was quantified by counting when the majority of progeny had at least reached the young adult stage (about 3 days after transfer).

### Sperm activation assay and imaging

To perform sperm activation assays, spermatids were dissected from adult males that were shifted to 25°C during the embryo stage, and isolated prior to sexual maturity (about 48 h post L1). Dissection was performed directly on glass slides in sperm medium (50 mM HEPES pH 7.8, 50 mM NaCl, 25 mM KCl, 5 mM CaCl_2_, and 1 mM MgSO_4_) supplemented with 20 μg/mL pronase E (Millipore Sigma). For the characterization of sperm morphology, sperm were imaged 30 min after the addition of pronase E. Individual sperm were manually categorized into two types: spermatids with normal pseudopods or spermatids with irregular or no pseudopods (Shakes and Ward, 1989). For Figure 5E, Z stacks were imaged in 10 sec intervals for 30 min and a representative in-focus stack was chosen at every 3 min interval. To characterize sperm activation dynamics, sperm were individually followed across 10 sec intervals for 30 min and the different stages of sperm activation were designated into four categories based on these morphological changes: 1) undifferentiated spermatid, 2) intermediate spindles characterized by the presence of spike growth, 3) growing or motile pseudopod by the presence of a pseudopod, and 4) immobile sperm when little movement was observed either in the sperm body or pseudopod for longer than 30 sec. Statistical significance was assessed using Student’s t-test

## QUANTITATIVE AND STATISTICAL ANALYSIS

Unless otherwise stated, all quantitative analyses are shown as mean with standard deviation represented as error bars. For qRT-PCR, fertility and mating assays, and western blot, at least 2 independent experiments were performed; one representative biological replicate is shown.

### Small RNA-seq analysis

Raw small RNA-seq reads were trimmed for Illumina adapters and quality (SLIDING WINDOW: 4:25) using Trimmomatic 0.39 (Bolger et al., 2014). Trimmed reads were then filtered using bbmap 38.23 (http://jgi.doe.gov/data-and-tools/bb-tools) to retain reads that were 15–30 nt in length. These filtered reads were aligned to the *C. elegans* WBcel235 (Cunningham et al., 2018) reference genome using Bowtie 1.1.1 (Langmead et al., 2009) with parameters -v 0 -k 5 –best –strata –tryhard. Quality control of raw and aligned reads was performed using FastQC 0.11.7 (http://www.bioinformatics.babraham.ac.uk/projects/fastqc/), SAMtools 1.9 (Li et al., 2009) and in-house Python and R scripts. Mapped reads were assigned to genomic features using featureCounts from Subread 1.6.3 (Liao et al., 2014), taking into account overlapping and multi-mapping reads (-O -M). Raw counts were normalized within DESeq2 1.26.0 (Love et al., 2014) and principal component analysis (PCA) was performed using the regularized log transform of normalized counts within DESeq2 (Figure S5C).

To identify differentially expressed genes, DESeq2 was applied to piRNAs on chromosome IV. In this study (method 1), we define significant and differentially expressed genes as having an absolute value of log_2_(fold-change) ≥ 0.26 and FDR of ≤ 0.05 (Benjamini-Hochberg). Contrasts between mutant and wildtype were designed without independent filtering.

For motif discovery, nucleotide sequences were extracted from the reference genome with 60 nt upstream of each piRNA and submitted to the MEME suite 5.1.1 (Bailey et al., 2009). Results from MEME were used to generate the Logo plot with the median position of the C-nucleotide of the identified motif, the number of piRNAs that share the motif, and the associated E-value.

A second, independent small RNA-seq analysis workflow (described in Figure S2) was implemented to validate our results. Results produced from this analysis are provided in Figure S3. 16–30 nt small RNA sequences were parsed from adapters and reads with >3 nt falling below a quality score of Q30 were discarded. Reads were mapped to the *C. elegans* WS230 (Stein et al., 2001) reference genome using CASHX v. 2.3 (Fahlgren et al., 2009) allowing for 0 mismatches. Custom Perl, Awk, and R scripts were used to count features and to generate PCA and size distribution plots. Multi-mapping reads were assigned proportionally to each possible locus. Differential expression analysis was done using DESeq2 v. 1.18.1 (Love et al., 2014). A reporting threshold was set at an absolute value of log_2_(fold-change) ≥ 0.26 and a Benjamini-Hochberg-corrected p ≤ 0.20.

### ChIP-seq analysis

De-multiplexed raw ChIP-seq data in FASTQ format were trimmed for adapters and sequencing quality score > Q25 using Trim Galore! 0.5.0 (http://www.bioinformatics.babraham.ac.uk/projects/trim_galore/) and aligned to *C. elegans* reference genome WBcel235 (Cunningham et al., 2018) using Bowtie2 2.3.4.2 (Langmead and Salzberg, 2012) with default parameters. Post-alignment filtering was then performed to remove PCR duplicates using the MarkDuplicates utility within Picard 2.22.1 (http://broadinstitute.github.io/picard/). In addition, SAMtools 1.9 was applied to remove unmapped reads and reads that mapped with MAPQ 30 but were not of primary alignment or failed sequence platform quality checks (SAMtools -F 1804 -q 30) (Li et al., 2009).

To identify and visualize binding sites and peaks for SNPC-1.3 ChIP-seq, filtered SNPC-1.3 ChIP-seq reads were extended to 200 bp to account for the average length of ChIP fragments. We then partitioned the genome into consecutive, non-overlapping 1 kb bins and calculated read coverage, normalized by sequencing depth of each library, based on the total read count in each bin. Bins with read coverages in the IP sample that fell below the median read coverage of piRNA-depleted bins on chromosome IV in the relevant input control were excluded from further analysis. Bins containing only male, female, and non-germline enriched piRNAs (as defined by small RNA-seq analysis) were then extracted to generate binding profiles and heatmaps. For this, the bamCompare tool in deepTools 3.3.1 (Ramírez et al., 2016) was used to calculate the ratio between read coverage of each ChIP sample and input control (--scaleFactorsMethod None --normalizeUsing CPM --operation ratio --binSize 50 --ignoreForNormalization MtDNA --extendReads 200). The ENCODE ce11 blacklist was also supplied (https://github.com/Boyle-Lab/Blacklist/tree/master/lists). The bamCompare output was then used in deepTools computeMatrix to calculate scores for plotting profiles and heatmaps with deepTools plotProfile and plotHeatmap.

TRA-1 ChIP-seq peaks were called by callpeak within MACS 2.1.2 (Zhang et al., 2008) (--pvalue 0.05) with filtered TRA-1 ChIP-seq reads and relevant input controls. TRA-1 signal tracks were generated by calculating fold enrichment from read count-normalized genome-wide pileup and lambda track outputs by callpeak (bdgcmp in MACS2). The ENCODE ce 11 blacklist was supplied in this analysis (https://github.com/Boyle-Lab/Blacklist/tree/master/lists). The bamCompare tool in deepTools 3.3.1 (Ramírez et al., 2016) was used to quantify read coverage of each ChIP sample and input control.

Reproducibility between SNPC-1.3 and TRA-1 ChIP-seq replicates (Figure S4C, S5E) was assessed by applying deepTools bamCompare, as described above, and deepTools plotCorrelation to depict pairwise correlations between replicates and compute the Pearson correlation coefficient.

## DATA AND SOFTWARE AVAILABILITY

The mass spectrometry, small RNA-seq, and ChIP-seq data have been deposited in NCBI under GEO accession number: GSE152831. Processed data and scripts used for analysis are available at https://github.com/starostikm/SNPC-1.3.

## AUTHOR CONTRIBUTIONS

Conceptualization of the study: CPC, RJT, and JKK; experimental design: CPC, RJT, and JKK; mass spectrometry analysis: JJM and JRY; ChIP library preparation and sequencing: SF and SEJ; RNA library preparation and sequencing: BEM and TAM; all other experiments: EX, MAH, RJT, CPC; computational analysis: MRS, CPC, TAM, MCS; CPC, RJT, MRS, and JKK wrote the manuscript.

## ACKNOWLEDGEMENTS

We thank Himani Galagali and Natasha Weiser for helpful comments on the manuscript. We thank members of the Kim Lab (Amelia Alessi, Mindy Clark, Gregory Fuller, Jessica Kirshner, Alex Rittenhouse, and Darius Mostaghimi), Tatjana Trcek, Angela Andersen, Aurelia Mapps, and Jacqueline Tay for helpful suggestions. Computational resources were provided by the Maryland Advanced Research Computing Center (MARCC). Some strains were provided by the *Caenorhabditis* Genetics Center, which is funded by the NIH Office of Research Infra-structure Programs (P40 OD010440). This work was supported by grants from the NSF DGE-1746891 (to R.J.T.), NIH R35 GM130272 (to S.E.J); NIH R35 GM119775 (to T.A.M.); and NIH R01 GM129301 and NIH R01 GM118875 (to J.K.K.).

## COMPETING INTERESTS

The authors declare no competing interests.

## SUPPLEMENTAL FIGURE LEGENDS

**Figure S1:**
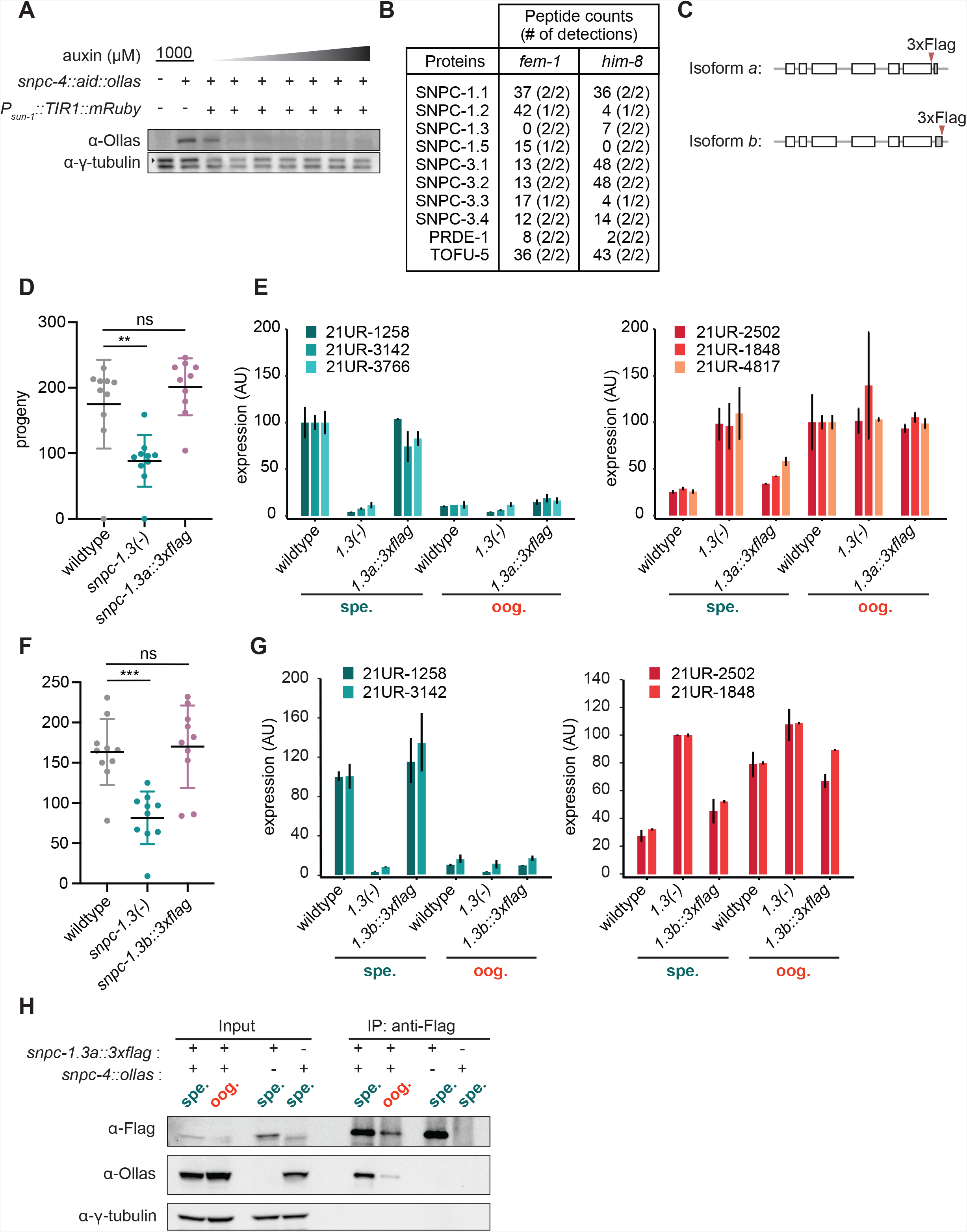
Related to Figure 1. Validation of strains and mass spectrometry. (A) SNPC-4::AID is substantially degraded at 250 μM auxin in the germline. Western blot of SNPC-4::AID::Ollas in worms placed on various auxin concentrations (0, 25, 50, 100, 250, 500, 1000 μM) for 1 h. The germline promoter *Psun-1* was used to drive expression of the *A. thaliana* TIR1. **γ**-tubulin is the loading control. (B) Table showing the peptide counts of SNAPc homologs and piRNA biogenesis proteins identified from mass spectrometry of immunopurified SNPC-4::3xFlag from *fem-1(-)* and *him-8(-)* strains. (C) *snpc-1.3a::3xflag* strain has wild-type fertility at 25°C. Black bars indicate mean ± SD of n=10 worms (wildtype versus *snpc-1.3(-)* mutant **p ≤ 0.005, Welch’s t-test). (D) *snpc-1.3a::3xflag* strain has wild-type levels of male and female piRNAs during spermatogenesis and oogenesis. *1.3(-)* denote*s snpc-1.3(-)*. *1.3a::3xflag* denotes *snpc-1.3a::3xflag.* Error bars indicate ± SD from two technical replicates. (E) s*npc-1.3b::3xflag* strain has wild-type fertility at 25°C. Black bars indicate mean ± SD of n=10 worms (wildtype versus *snpc-1.3(-)* mutant ***p ≤ 0.0001, Welch’s t-test). (F) *snpc-1.3b::3xflag* strain has wild-type levels of male and female piRNAs during spermatogenesis and oogenesis. *1.3(-)* denote*s snpc-1.3(-). 1.3b::3xflag* denotes *snpc-1.3b::3xflag.* Error bars indicate ± SD from two technical replicates. (G) SNPC-1.3 interacts with SNPC-4. Reciprocal immunoprecipitation of Figure 1E. Anti-Flag immunoprecipitation of SNPC-1.3::3xFlag and Western blot of SNPC-4::Ollas during spermatogenesis and oogenesis. **γ**-tubulin is the loading control.

**Figure S2:**
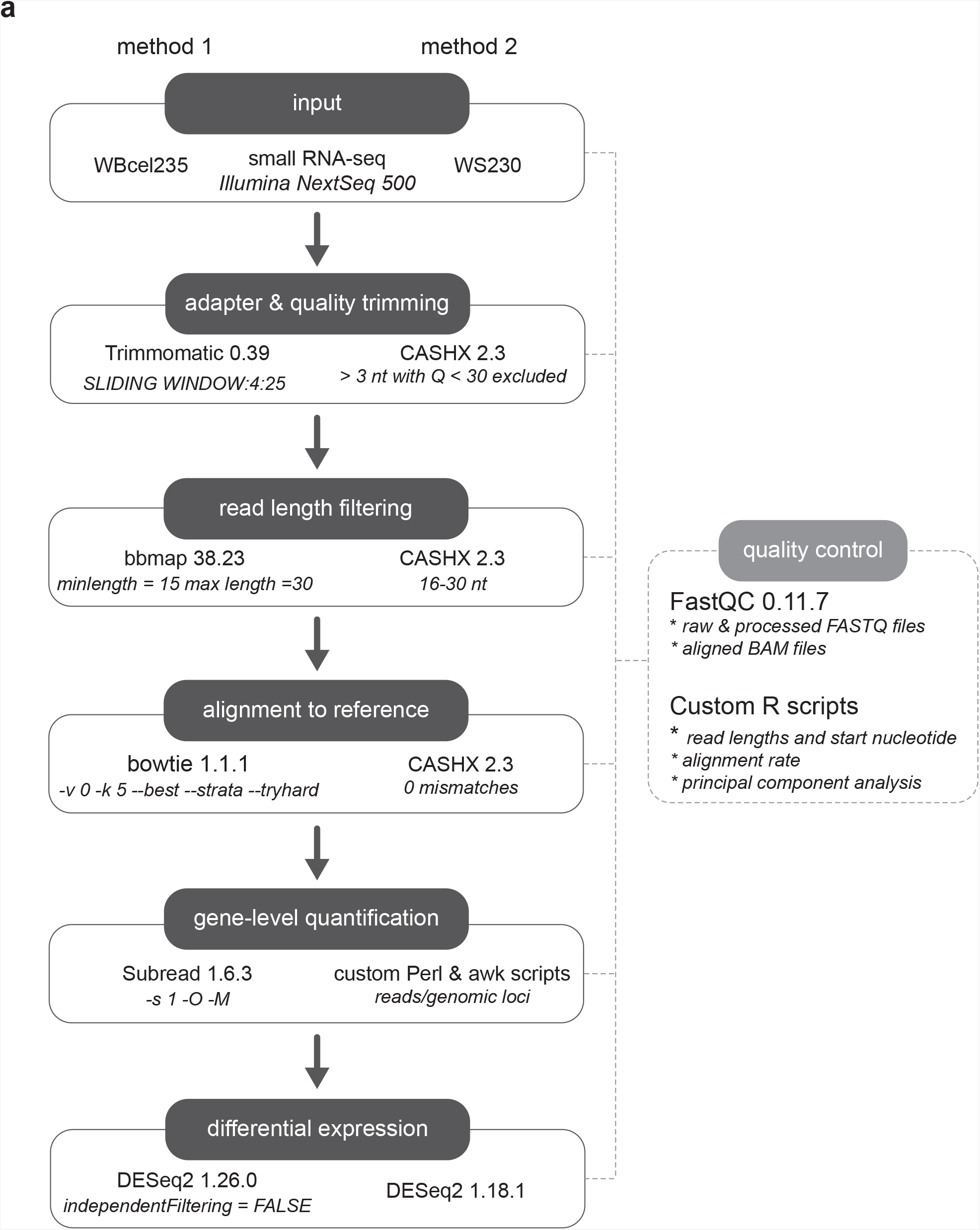
Related to Figure 2. Small RNA-seq analysis pipeline. Two independent workflows (method 1 and method 2) were applied for differential expression analysis.

**Figure S3:**
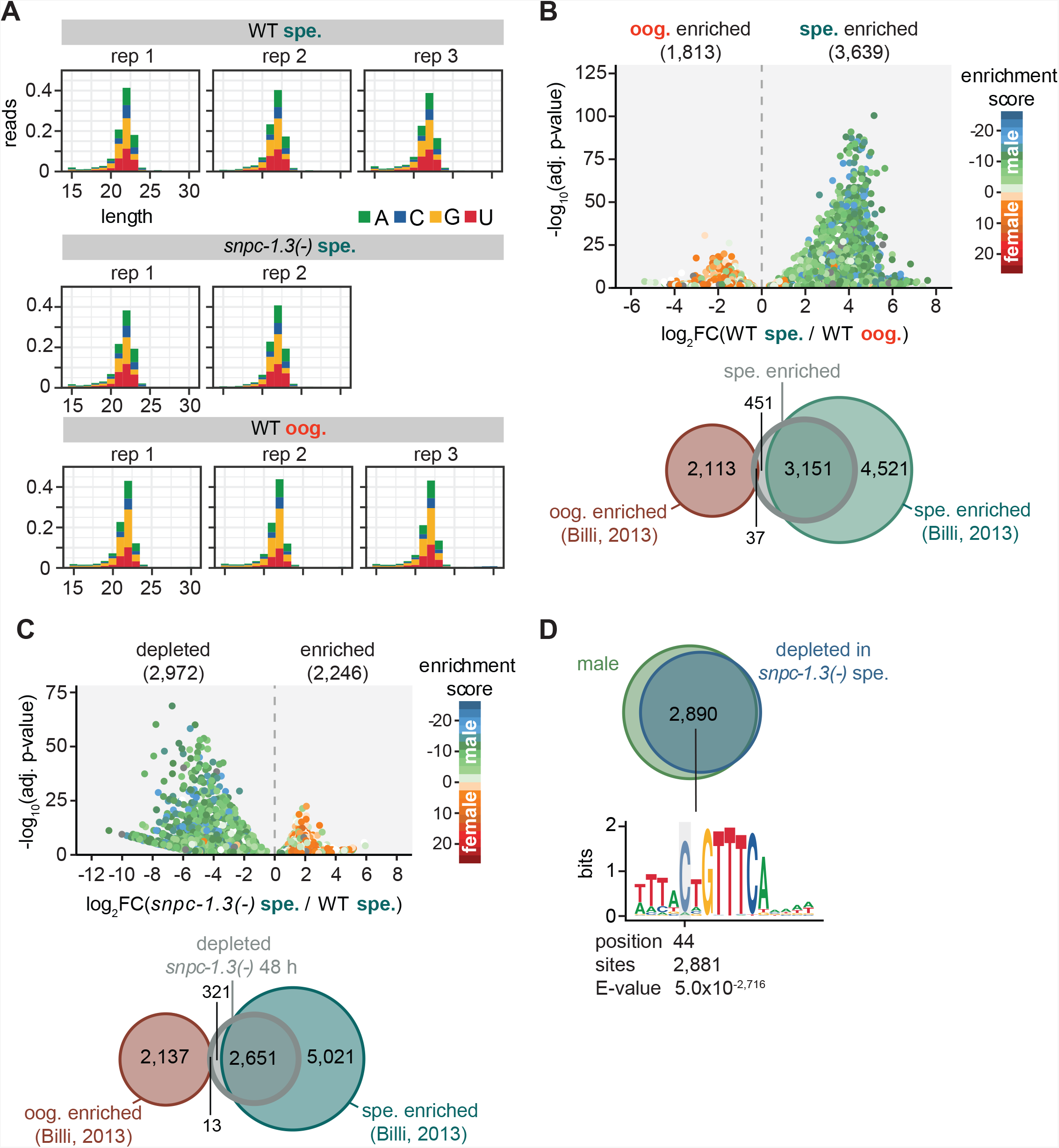
Related to Figure 2. Quality control of small RNA-seq and validation analysis. (A) Mapped reads distributed by read length and 5’ nucleotide identity (B) (Top) Volcano plot showing differential piRNA expression between spermatogenesis and oogenesis in wild-type worms. piRNAs are colored according to male and female enrichment scores from Billi et al., 2013. Analysis shown is from a second, independent small RNA-seq analysis workflow (method 2). (Bottom) Overlap of male piRNAs in wildtype at 48 h with oogenesis- and spermatogenesis-enriched piRNAs defined in Billi et al., 2013. (C) (Top) Volcano plot showing differential piRNA expression between *snpc-1.3(-)* mutants versus wildtype during spermatogenesis. piRNAs are colored by enrichment scores of male and female piRNAs defined in Billi et al., 2013. Analysis shown is from a second, independent small RNA-seq analysis workflow (method 2). (Bottom) Overlap of SNPC-1.3-dependent piRNAs with previously identified spermatogenesis- and oogenesis-enriched piRNAs (Billi et al., 2013). (D) Most SNPC-1.3-dependent piRNAs overlap with male piRNAs (as defined in S3B) (top) and are enriched for the upstream 8 nt core motif showing a bias for C at the 5’ position (bottom).

**Figure S4:**
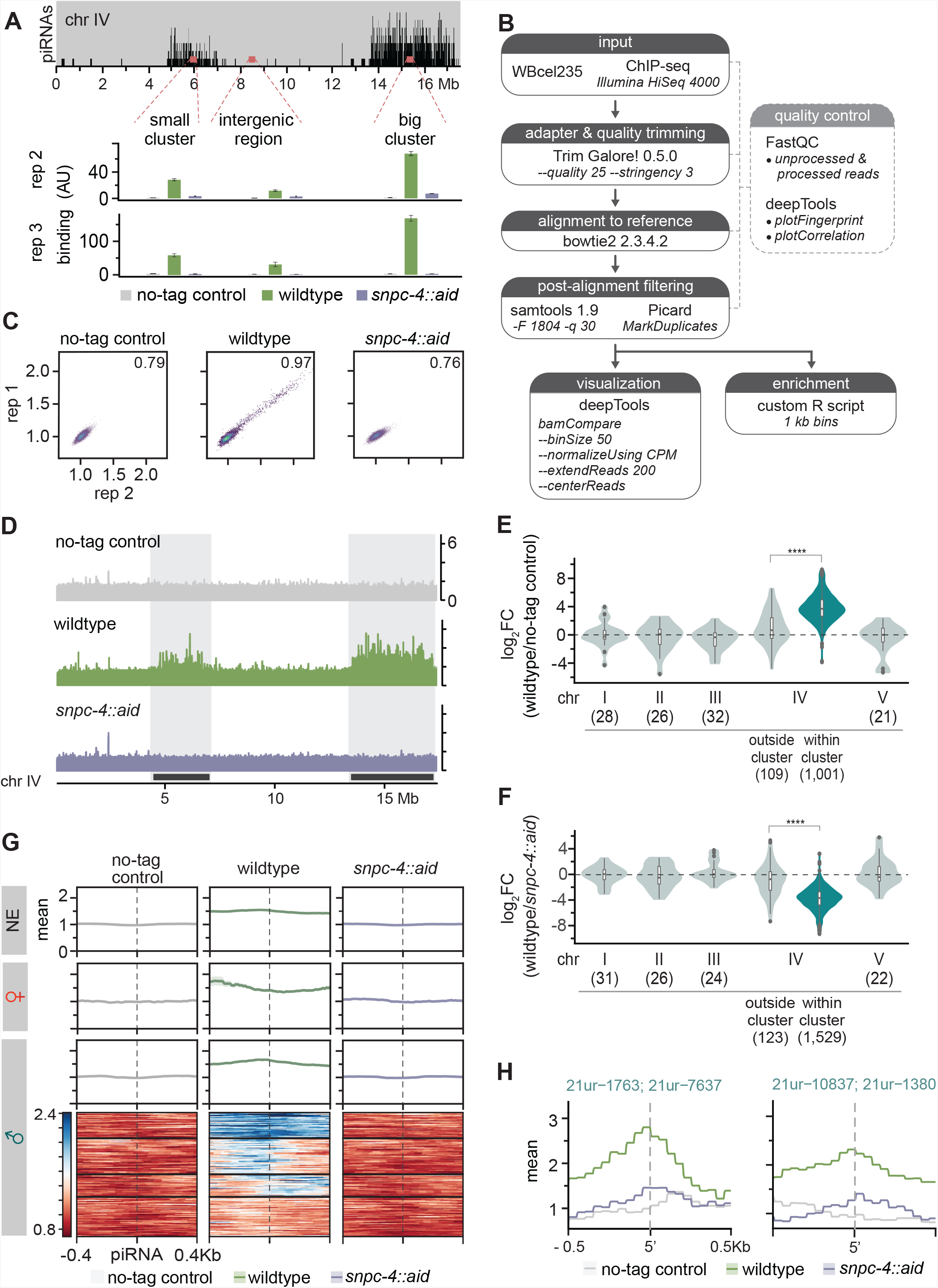
Related to Figure 3. SNPC-1.3 ChIP-seq pipeline and quality control and biological replicates for SNPC-1.3 ChIP. (A) SNPC-1.3 binding at the piRNA clusters requires SNPC-4. Two biological replicates of the experiment shown in Figure 3A. Top panel depicts the density of piRNAs on chromosome IV, showing piRNAs are predominantly found in a small (4.5–7 Mb) and big (13.5–17.2 Mb) cluster. SNPC-1.3::3xFlag binding normalized to input (mean ± SD of two technical replicates) on chromosome IV by ChIP-qPCR in a no-tag control, the strain expressing SNPC-4::3xFlag (wildtype), and in the strain expressing SNPC-4::3xFlag::AID, which undergoes TIR-1-mediated degradation by addition of auxin (*snpc-4::aid*). These labels (no-tag, wild-type, and *snpc-4::aid*) apply throughout Figure S4. (B) SNPC-1.3 ChIP-seq analysis workflow. (C) Pairwise Pearson correlations between SNPC-1.3 ChIP-seq biological replicates. (D) Biological replicate of Figure 3B. The locations of the two piRNAs clusters are highlighted. (E) SNPC-1.3 binding is enriched at piRNA clusters on chromosome IV. Biological replicate of Figure 3C. Regions within piRNA clusters are enriched for SNPC-1.3 binding compared to regions outside of the piRNA clusters on chromosome IV (**** p≤ 0.0001, Wilcoxon rank sum test). The number of bins analyzed is listed in parentheses. (F) SNPC-1.3 enrichment at piRNA clusters is dependent on SNPC-4. Biological replicate of Figure 3D. SNPC-1.3-bound regions within piRNA clusters are depleted compared to regions outside of the piRNA clusters on chromosome IV upon loss of SNPC-4 (**** p≤ 0.0001, Wilcoxon rank sum test). The number of bins analyzed is listed in parentheses. (G) Biological replicate of enrichment profiles shown in Figure 3E. Distribution of SNPC-1.3 reads (mean density ± standard error) around the 5’ nucleotide of mature piRNAs at the piRNA clusters. To resolve SNPC-1.3 binding between male and female piRNAs despite the high density of piRNAs, we selected 1 kb bins with all male (135), female (20), or non-germline enriched (337) piRNAs. (H) Examples of SNPC-1.3 binding at two regions containing two male piRNA loci. Regions are anchored on the 5’ nucleotide of each mature male piRNA and show mean read density ± standard error.

**Figure S5:**
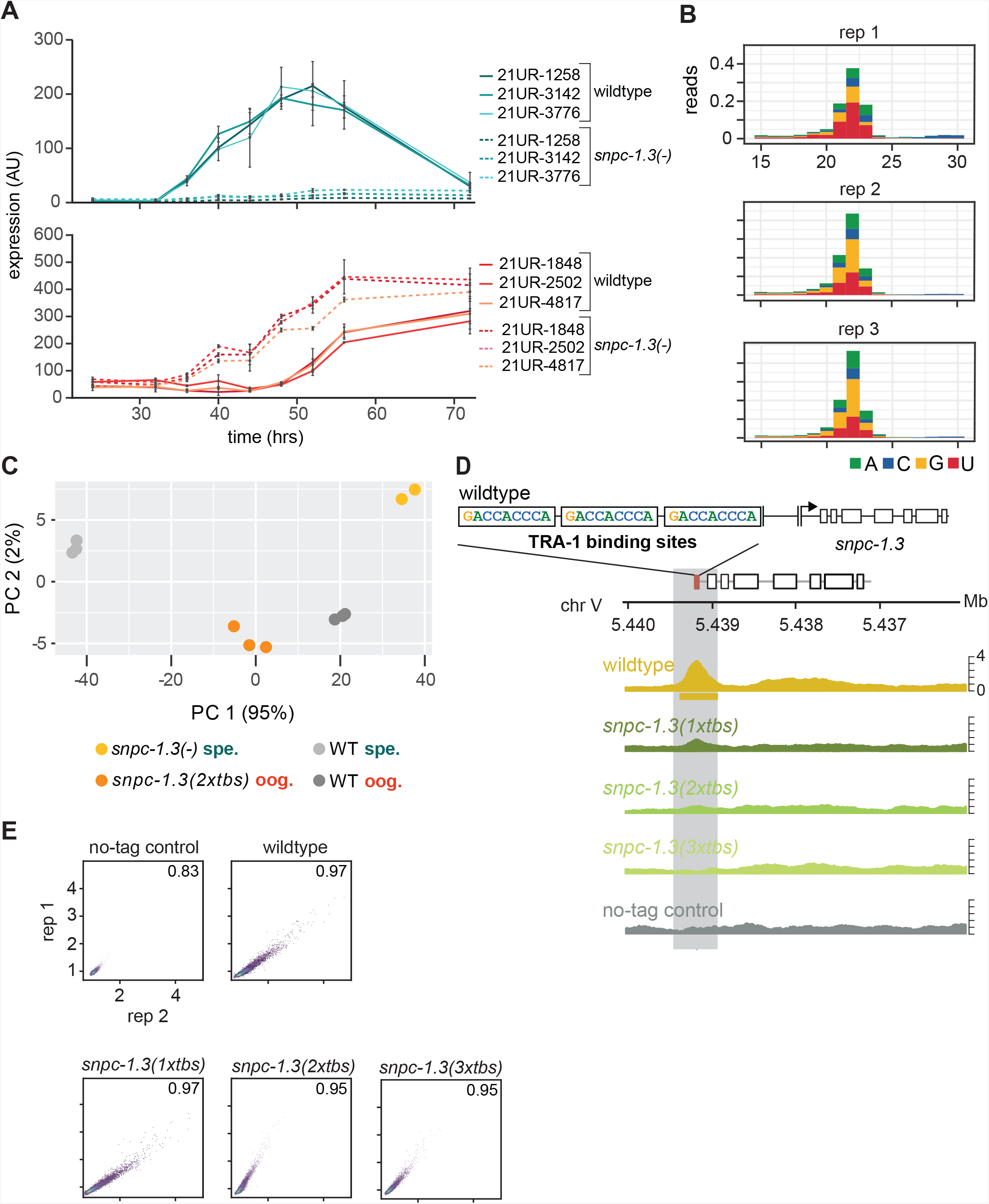
Related to Figure 4. TRA-1 regulation of *snpc-1.3* across hermaphrodite development. (A) Male and female piRNA levels peak during spermatogenesis and oogenesis, respectively. Male piRNA levels are severely impaired but female piRNA expression is upregulated during hermaphrodite development in *snpc-1.3* mutants. Taqman qPCR and quantification of male and female piRNAs across hermaphrodite development. Error bars: ± SD of two technical replicates. (B) Mapped reads distributed by read length and 5’ nucleotide identity of 3 biological replicates. (C) Principal component (PC) analysis of piRNA expression based on rlog transformation of normalized counts. The scatter plot depicts the first two principal components. Percentage on each axis represents the percentage of variation accounted for by each principal component. (D) TRA-1 binds to the *snpc-1.3* promoter. Biological replicate of Figure 4B. Schematic of the three TRA-1 binding sites upstream of the *snpc-1.3* locus in wildtype (top). (Bottom) TRA-1 binding is progressively reduced with the increase in number of TRA-1 binding sites mutated. (E) Pairwise Pearson correlations between TRA-1 ChIP-seq biological replicates.

**Figure S6:**
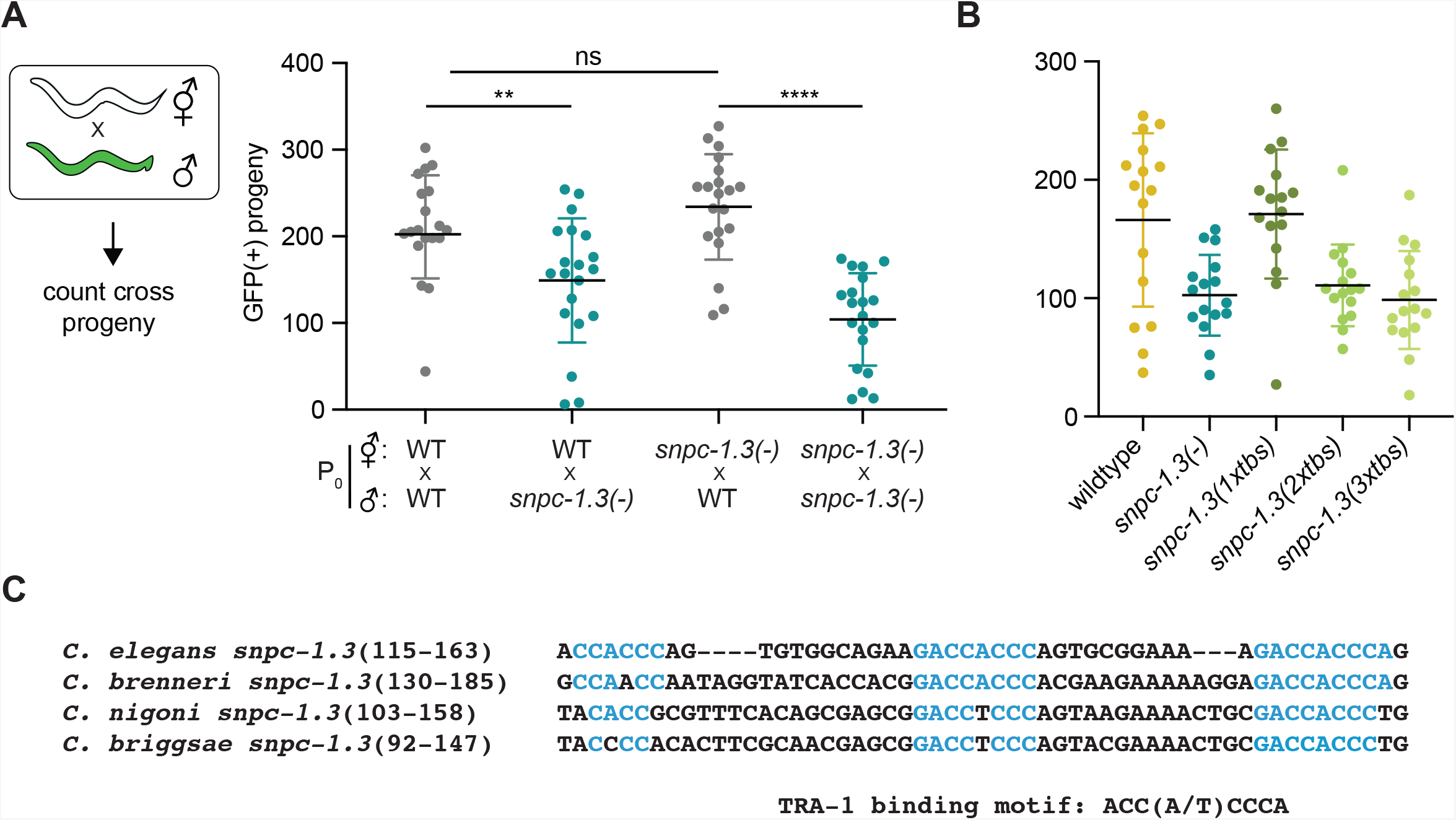
Related to Figure 5. SNPC-1.3 is critical for male fertility. (A) *snpc-1.3* is required in males, but not females, to promote fertility. (Left) Diagram illustrates crosses between strains for mating assays. Green worms represent *col-19::GFP* males. (Right) *snpc-1.3(-); col-19::GFP* males show diminished ability to rescue wild-type or *snpc-1.3(-)* hermaphrodites when compared to *col-19::GFP* males. Circles correspond to the brood size of viable progeny from each mating (n=16). Black bars indicate mean ± SD. Statistical significance was assessed using Welch’s t-test (ns: not significant;**p ≤ 0.005; ****p ≤ 0.0005). (B) *snpc-1.3(2xtbs)* and *snpc-1.3(3xtbs)* mutant hermaphrodites have decreased fertility at 25°C. Black bars indicate mean ± SD (n = 16). Statistical significance was assessed using Welch’s t-test (****p ≤ 0.0001). (C) Multiple TRA-1 binding sites are conserved in *C. elegans, C. brenneri, C. briggsae,* and *C. nigoni.* Sequence alignment of *snpc-1.3* homologs in other nematode species. Blue indicates sequences in TRA-1 binding motifs.

---

**Table S1: Differential expression of piRNAs in wild-type worms during spermatogenesis and oogenesis. Related to Figure 2.**

**Table S2: Differential expression of piRNAs in wildtype and***snpc-1.3(-)* **mutants during spermatogenesis. Related to Figure 2.**

**Table S3: Differential expression of piRNAs in wildtype and***snpc-1.3(2xtbs)* mutants during oogenesis. Related to Figure 4.

**Table S4. List of strains, guides, repair templates, and oligos used in this study**

## Notes

### Competing Interest Statement

The authors have declared no competing interest.

